# Discovering functionally important sites in proteins

**DOI:** 10.1101/2022.07.14.500015

**Authors:** Matteo Cagiada, Sandro Bottaro, Søren Lindemose, Signe M. Schenstrøm, Amelie Stein, Rasmus Hartmann-Petersen, Kresten Lindorff-Larsen

## Abstract

Proteins play important roles in biology, biotechnology and pharmacology, and missense variants are a common cause of disease. Discovering functionally important sites in proteins is a central but difficult problem because of the lack of large, systematic data sets. Sequence conservation can highlight residues that are functionally important but is often convoluted with a signal for preserving structural stability. We here present a machine learning method to predict functional sites by combining statistical models for protein sequences with biophysical models of stability. We train the model using multiplexed experimental data on variant effects and validate it broadly. We show how the model can be used to discover active sites, as well as regulatory and binding sites. We illustrate the utility of the model by prospective prediction and subsequent experimental validation on the functional consequences of missense variants in *HPRT1* which may cause Lesch-Nyhan syndrome, and pinpoint the molecular mechanisms by which they cause disease.

## Introduction

Proteins carry out most functions in a cell, generally through interactions with other molecules. These molecular interactions often involve specific sites or regions whose identification plays a fundamental role in understanding biology and disease. Progress has been made in the development of methods to identify some types of functional sites (del Sol Mesa et al., 2003; Lee et al., 2007; Radivojac et al., 2013; Kulmanov et al., 2018; Torng and Altman, 2019) and efforts have been made to understand the relationship between sequence variability, protein function, stability and the onset of diseases (Yue et al., 2005; Adzhubei et al., 2010; Kircher et al., 2014; Wagih et al., 2018; Livesey and Marsh, 2020). With the entry of accurate and large-scale protein structure prediction, structure-based methods for understanding biology are becoming even more important.

Analyses of large-scale mutagenesis studies have been used to probe the role of individual residues in the stability, abundance and function of a protein (Gray et al., 2017; Dunham and Beltrao, 2021; Høie et al., 2022). Since most proteins need to be folded to function, it may, however, be difficult to deconvolute the effects of amino acid substitutions on intrinsic function from their effects on stability and cellular abundance (Li and Lehner, 2020). We note here that we use the term ‘function’ and ‘functional sites’ in a relatively general sense since our goal is to examine protein function broadly. In a few, favourable cases, multiplexed assays of variant effects (MAVEs) have been used to probe the consequence of almost all individual substitutions using both a functional readout and a readout that probes cellular abundance. Analysis of such data have been used to shed light on the molecular mechanisms underlying perturbed function, and more specifically to pinpoint which functional properties an amino acid residue contributes to (Jepsen et al., 2020; Cagiada et al., 2021; Chiasson et al., 2020; Faure et al., 2022). For example, variants that lose function together with loss of abundance are likely to be caused by perturbations to the overall protein fold and stability, whereas variants that lose function while retaining wild-type-like abundance in the cell are likely to be caused by perturbing sites that directly play a role in function (Jepsen et al., 2020; Cagiada et al., 2021; Faure et al., 2022). Alternative to such multi-readout experiments, global analyses of large datasets of multi-mutant variants can be used to deconvolute effects on for example stability and binding (Otwinowski, 2018).

In a recent analysis of abundance and activity assays (Cagiada et al., 2021) we showed that approximately half of the single-point variants that show loss-of-function do so together with loss of protein abundance. This result suggests that if one uses a functional readout to detect residues that are directly involved in function, then about half of the variants detected as important are so simply because they cause lowered abundance. This in turn makes it difficult to separate residues that are directly involved in function (for example in catalysis, binding and signalling) from those positions that are conserved mostly due to structural constraints (Echave and Wilke, 2017).

The other half of loss-of-function substitutions, instead, mostly affects the protein via for example perturbing interactions with substrates or binding partners rather than stability (Cagiada et al., 2021; Nielsen et al., 2021). We call this class of substitutions ‘stable but inactive’ (SBI) variants emphasising their tight and direct involvement in protein function. Pinpointing and predicting which variants are SBI is important to understand how amino acid substitutions might affect protein functions, why they possibly cause disease, and to ultimately aid the development of personalised therapeutic treatments. A very practical utility of the detection of SBI variants is that they enable the identification of amino acid residues that play a direct role in function. Indeed, much of our biochemical understanding of how proteins function relies of protein engineering studies where the effects of amino acid substitutions on various biochemical readouts are probed. Thus, positions where most substitutions affect function, but not structural stability, are often found in functional sites (Cagiada et al., 2021; Chiasson et al., 2020; Faure et al., 2022).

As an alternative to experimental measurements, computational predictions can offer a faster, cheaper and more scalable approach to pinpoint positions that are important for function. Computational prediction of SBI variants, however, is not trivial. Many existing computational protocols are based on evaluating sequence conservation to find positions and variants that cause loss of function (Lichtarge et al., 1996; Kumar et al., 2009; Choi and Chan, 2015; Riesselman et al., 2018; Laine et al., 2019; Torng and Altman, 2019). This strategy alone, however, may not be sufficient to identify residues that are conserved due to direct functional roles because sequence evolution is subject to both functional and structural constraints, and thus the different signals cannot be easily disentangled. One possible strategy to overcome this problem is to combine evolutionary data with analyses that report directly on the effects of substitutions on protein stability (Cheng et al., 2005; Wang et al., 2008; Capra et al., 2009; Cagiada et al., 2021).

Here, we aim to create a robust and easy-to-use prediction method to identify functional residues in proteins and provide insights into the biochemical role of these residues in the target protein. To avoid biases from annotations of functional sites, we use data generated by MAVEs that report on the effects of a wide range of substitutions on both function and abundance to train a supervised machine learning model. As input to the model we use a combination of sequence conservation measures, free energy changes accompanying substitutions, and physicochemical properties. We begin by showing how our model can capture a wide range of functional sites that include those in active sites, but also distributed throughout the protein structure including potential allosteric and regulatory sites. We then show how we achieve good accuracy in pinpointing functional amino acids in different validation scenarios. Across several proteins we find that roughly one in ten of the positions are functionally relevant and conserved for reasons different than structural stability. Having validated the model, we use it to provide examples of the kinds of insight that it may provide including identifying catalytic sites, regions that interact with substrates, and interfaces in complexes. Finally, we performed prospective predictions of the mechanisms of disease variants in HPRT1 (encoding the protein Hypoxanthine-Guanine Phosphoribosyltransferase 1, HPRT1) and validate these by measuring variant effects on function and abundance. The code for our model is freely available and we also provide access to it via a web implementation.

## Results

### Identification of functional sites via analyses of stability and conservation

Previously we have used evolutionary analyses combined with stability calculations to predict variant effects on protein function and stability and showed how these measures correlate with changes in cellular abundance or function (Cagiada et al., 2021; Høie et al., 2022). Here, we build on these ideas to construct a model to identify functional sites in proteins via the identification of SBI variants (Fig.1). Before detailing the model, we first provide an intuitive description of the basic idea. Our goal is to identify positions in proteins that play some role in protein function and regulation; these may for example include active sites in enzymes, but also ‘second-shell’ residues around the active site, protein-protein interfaces, allosteric and regulatory sites, or residues involved in recognizing ligands and substrates. Experimentally, these residues may be identified since amino acid substitutions at these sites might cause loss of ‘function’ (in some readout). Many variants, however, cause loss of function via loss of stability and cellular abundance, and we remove such indirect effects by requiring that the variants have close to wild-type-like abundance (i.e. the variants belong to the SBI class). Computationally, we can identify these variants by calculations of effects on protein stability and conservation, thus finding positions that are conserved during evolution, but not due to a role in protein stability. We note two effects of these choices. First, the broad definition means that we can assign a functional role to a relatively large number of residues e.g. well beyond active sites in enzymes. Second, some residues will have a direct role in function, but also be important for protein stability; our analysis will miss those residues, but as shown below, our results suggest that most functional sites do not fall in this class.

**Figure 1.**
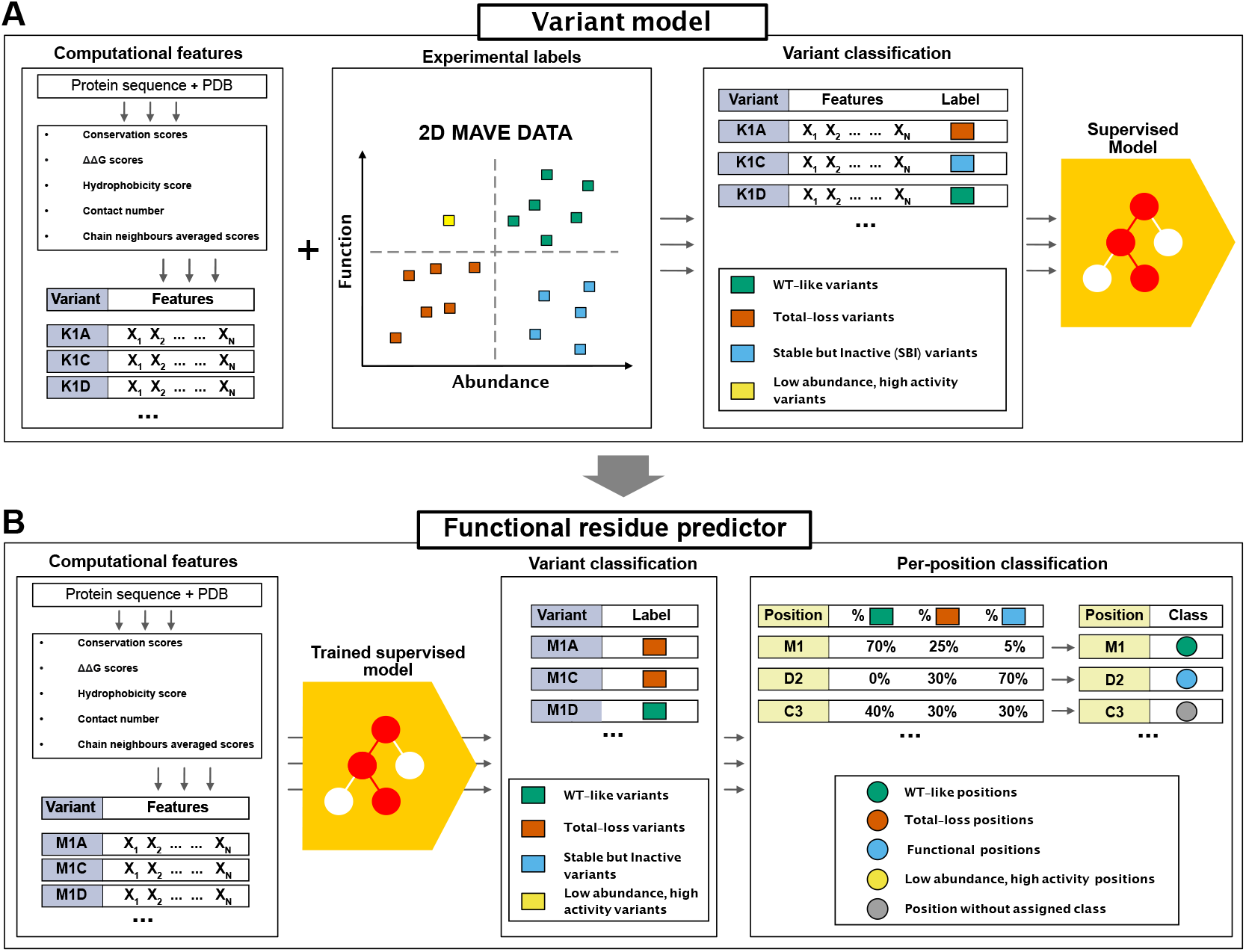
Graphical summary of our model to identify functionally important residues. (A) Using protein sequence and structure as input, we extract a number of features to characterise each variant. For each protein in our training set we extract all variants with MAVE measurements of abundance and function; this data is then combined with the structure and sequence features to train a gradient boosting classifier that assigns the variants to one of the four output classes. (B) The trained model takes the structure- and sequence-based features as input to classify all variants in to one of four classes. We also assign a class to each position/residue if half of the variants at that position are found in that class; remaining positions are not classified.

Based on the ideas above, we collected the results from two complementary types of MAVEs that respectively probe cellular-abundance and functional effects for three different proteins: NUDT15 (Suiter et al., 2020), PTEN (Matreyek et al., 2018; Mighell et al., 2018) and CYP2C9 (Amorosi et al., 2021) for a total of 9945 variants at 923 positions. Based on the experimental abundance and function scores, we assigned each variant to one of the following four classes (Cagiada et al., 2021): WT-like (high abundance and high activity), total loss (low abundance and low activity), SBI (high abundance and low activity), or low abundance and high activity. We also selected input features for each variant. These were calculated from the three-dimensional structure of a protein and a multiple sequence alignment. More precisely, we included the following features: (i) the predicted change in thermodynamic protein stability (ΔΔ*G*) calculated using Rosetta (Park et al., 2016), (ii) the evolutionary sequence information scores (which we term ΔΔ*E*, by analogy with ΔΔ*G*) us-ing GEMME (Laine et al., 2019), (iii) the hydrophobicity of the amino acid (Monera et al., 1995), and (iv) the weighted contact number (Shih et al., 2012; Jack et al., 2016) (Fig.1A).

We first examined whether one of these features on its own would be sufficient, but did not find any that individually separates SBI variants from the remainder (Fig.S1). Thus, we used the experimental labels and the input features to train a gradient boosting classifier (Fig.1A and Fig.S2) that predicts abundance (high/low) and activity (high/low) for each variant (see Methods for further details on the feature choices). We determined model hyperparameters using a stratified cross-validation procedure, where each validation set contained 20% of the data. The resulting model has an average validation accuracy of 58% and a Matthews correlation coefficient of 0.57. On the entire training data the model classified 9540/9945 variants (95%) correctly, of which 1638/1819 (90%) are SBI variants. Having assigned effects of the individual variants, we then used this data to pinpoint the functional residues in the target proteins. To this end, we assigned a residue to a class if at least half of the substitutions at that position belonged to that class. In particular, we focused our attention on amino acids classified as ‘functional residues’, where 50% or more of their variants are stable but inactive (Fig.1B). We compared the functional positions identified from the MAVEs with the predictions from the model (Fig.2). For the three proteins included in the training we correctly identify 116 out of 127 residues. Accuracy and true positive rates are similar for the three proteins (Fig.2).

**Figure 2.**
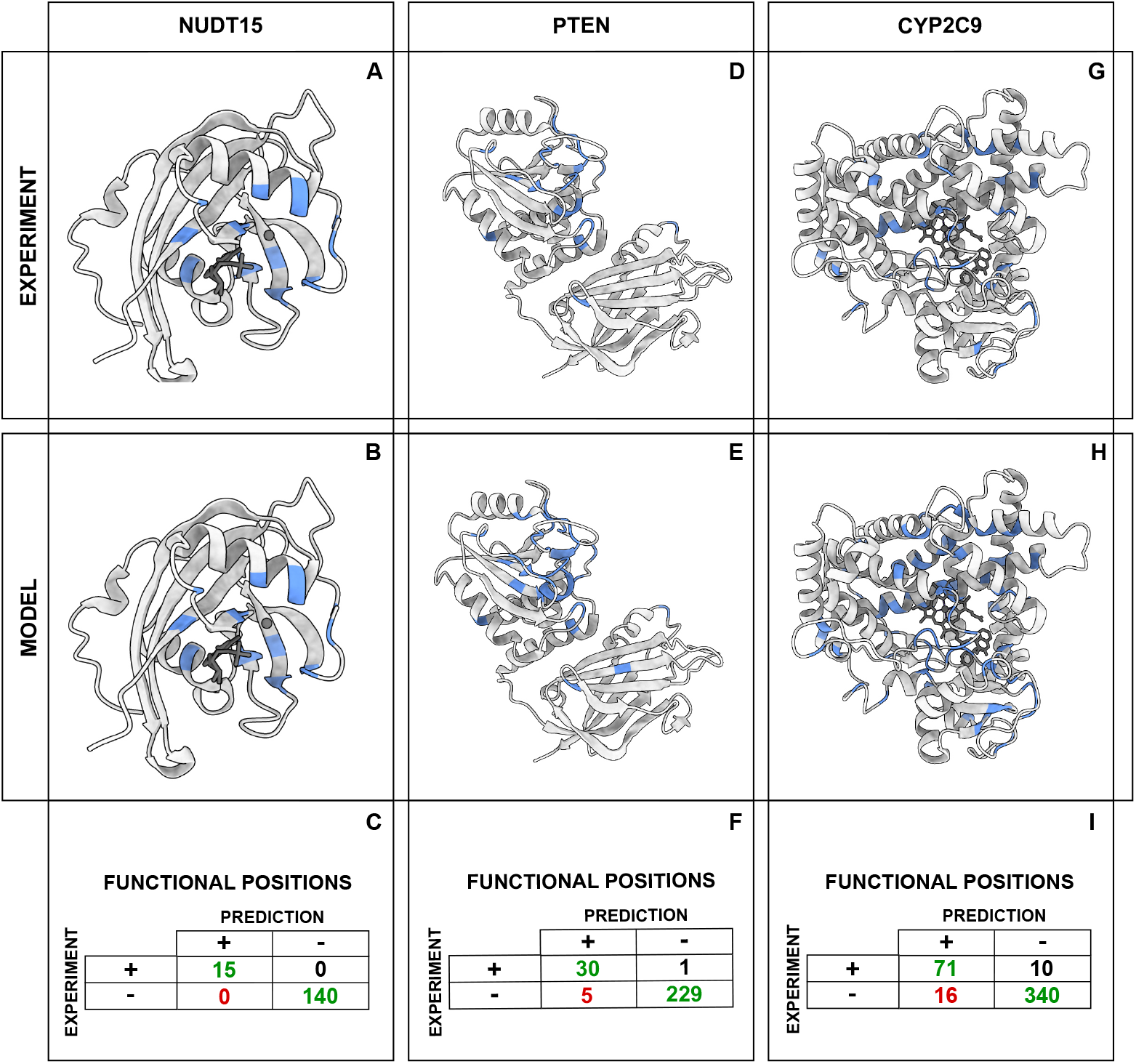
Finding functional residues in the training proteins. (A, D and G) Functional positions (in blue) identified using experiments. (B, E and H) Functional positions (in blue) identified using our model. (C, F, I) Statistics on the accuracy of residue level predictions of functional residues. For each of the three proteins we show the number of true positives (upper-left corner), true negatives (bottom-right corner), false positives (bottom-left corner) and false negatives (top-right).

After feature selection and training of the final model, we tested its performance on an independent dataset (GRB2 SH3 domain, Faure et al. (2022)) using as baseline a model using only cutoff values for ΔΔ*E* and ΔΔ*G* (Fig.S3A). We find that our model substantially outperforms the baseline model, especially in labelling functional residues. We also validated the performance of the model on a reduced variant training set (TableS2), and by training on two of the proteins and evaluating on the third (TableS3); in both cases we found reduced performance compared to the full model, but no evidence of substantial overfitting. We also examined other choices of features including both subsets of those used in our final model and a model with a wider set of features, and evaluated both by cross-validation and the independent dataset (Fig.S3B). The results show that the chosen feature set outperforms the other sets, including the model with a larger number of features.

### Validation and applications

Our model predicts sites and residues that play a broad range of functional roles. This, however, makes it more difficult to validate the model as most experiments only probe the role of a small number of sites. We therefore tested the method on diverse tasks and data.

### Validation using multiplexed and high-throughput data

First, we tested our model against a set of data that is relatively similar to the training data. Specifically, we applied it to two proteins (an SH3 domain from GRB2; Uniprot ID P62993 and the PDZ3 domain from PSD95; Uniprot ID P78352) that each have been assayed using two MAVEs that probe abundance and binding (Faure et al., 2022). A joint analysis of this data in turn enabled determining ΔΔ*G* for both folding and binding. We then used our model to predict variants and residues that are important for folding and function (here binding), and compared the results to the experiments (Fig.S5). For the SH3 domain, we find a good agreement between the predictions and experiments. Specifically, we find that variants and residues that are predicted to cause loss of function due to loss of abundance are generally found to have a large ΔΔ*G* for folding in the experiments (Fig.S5A,B). Similarly, the variants and residues that we predict to cause loss of function without loss of abundance, generally have a large ΔΔ*G* for binding, but are experimentally found not to affect folding. Note that our model did not use information about binding interactions, but simply discovers residues important for peptide binding by the fact that they are conserved, but not for stability reasons. A similar analysis of the PDZ domain showed a more complex picture, where we find a correlation (rather than separation) between the ΔΔ*G* for folding and binding, and that the predictions appear to divide the variants and residues into three categories with progressively greater ΔΔ*G* values (Fig.S5C,D). We note that for PDZ3 we found a limited number of homologs in Uniref30 for the PDZ3 domain, which might make the ΔΔ*E* values more uncertain.

We then analysed data from MAVEs on five different proteins to validate the performance of the variant classification step (Table1). In contrast to the data that were used to train the model, we here only had data for a single MAVE for each protein. Four of these experiments are functional assays and the last a cellular-abundance assay. Because these experiments only probe one dimension of our two-dimensional ‘function-and-abundance’ landscape, we simplified the output labels from the model from four to two classes. For the four MAVEs that probe function we combined the model’s output into either functional or inactive (independent of the predicted effect on abundance), and for the abundance-based MAVE we combine the predicted labels into high/low abundance (independent of the predicted effect on function). We compared these predictions to the data generated by the MAVEs which we also reduced to binary labels. On average we find an accuracy of 72% over all the variants and 68% when focusing on the variants labelled as SBI by our model (Table1).

We also compared our predictions with the results of a high-throughput experiment on alkaline phosphatase (PafA; Uniprot ID Q9KJX5, Markin et al. (2021)). In these experiments, the catalytic efficiency (*k*_cat_/*K*_M_) was measured for the wild-type protein as well as variants where each residue was changed into either glycine or valine. We first analyzed how well our model was able to distinguish between variants with different levels of activity, and found a lower median activity (*k*_cat_/*K*_M_) for variants classified as total-loss and SBI compared to WT-like variants (Fig.S6A,B). The experiments also revealed different effects of valine and glycine variants, and we analysed this observation in light of our predictions. For buried positions, we find that that variants at total-loss positions generally gave rise to lowered activity, in particular when substituting to glycine (Fig.S6C). At exposed positions, we generally predicted variants to have a smaller effect (Fig.S6D). While the experiments revealed that most variants did not cause global unfolding, a number of variants appeared to affect the *in vitro* translation that was used to produce the proteins (Markin et al., 2021). In particular, some variants resulted in less active protein when expressed at 37^◦^C compared to 23^◦^C. Using a ten-fold change as a cutoff (Markin et al., 2021), we find most (70%) variants that show this level of difference belong to the total-loss class (Fig.S6E,F), in line with the fact that these are predicted to cause folding defects.

Finally, we compared predictions from our model with results from two previously published models for detecting functional residues (Chelliah et al., 2004; Cheng et al., 2005). We selected 10 enzymes that had previously been studied and for which the percentage of true positives from the previous work was available. Using our model, we find 77 out of the 109 known functional sites (true positive rate of 0.71) (Fig.S4), which can be compared to values of 0.40 and 0.60 from the earlier models (Chelliah et al., 2004; Cheng et al., 2005).

### Predicting functional sites in enzymes and protein interactions

As an example of the kinds of insights one might gain on specific proteins, we applied the model to the Anti-sigma F factor (Uniprot: O32727, Fig.3A), for which previous studies have shown the importance of a number of sites (Campbell et al., 2002; Fu et al., 2019). Our model predicted 509 SBI variants (20% of the total number of variants) and we used these to label 21 positions as predicted functional sites. When comparing these to previous biochemical studies, we found reported functional roles for 17 of the 21 predicted positions (Campbell et al., 2002; Masuda et al., 2004). Of these 17 amino acids, nine are located in the proximity to the active site (Fig3B). His54, Gly55, Thr99, Gly107, Gly109 and Thr130 interact and stabilise the ADP/ATP in the active site, Asn50 is involved in the chelation of a Mg^2+^ ion, Glu46 acts as the catalytic base in the phosphorylation reaction and Arg105 stabilises the transition state in the phosphorylation reaction. In addition to these active site residues, we found another cluster of functional residues (Glu104, Thr49, Glu16, Ser45, Glu39 and Arg20) in the proximity to the binding site for the Anti-sigma F factor antagonist (Uniprot O32723, Fig.3B), while predicted functional positions Arg20 and Lys41 have been described as mediators of the interaction of the enzyme with the sigma factor. The roles of the remaining four positions that our model highlighted (Asn3, Asn15, Gly129 and Pro95) have, to our knowledge, not been analysed; they could either be false positives or residues with functional roles that have not yet been characterized. We observed that the functional sites in the Anti-sigma F factor could be grouped in two structurally compact clusters (Fig.S7). The first cluster includes all of the residues that have a role in the catalytic process, while the second cluster contains positions in the interaction network with Anti-sigma F factor antagonist and the sigma-factor.

**Figure 3.**
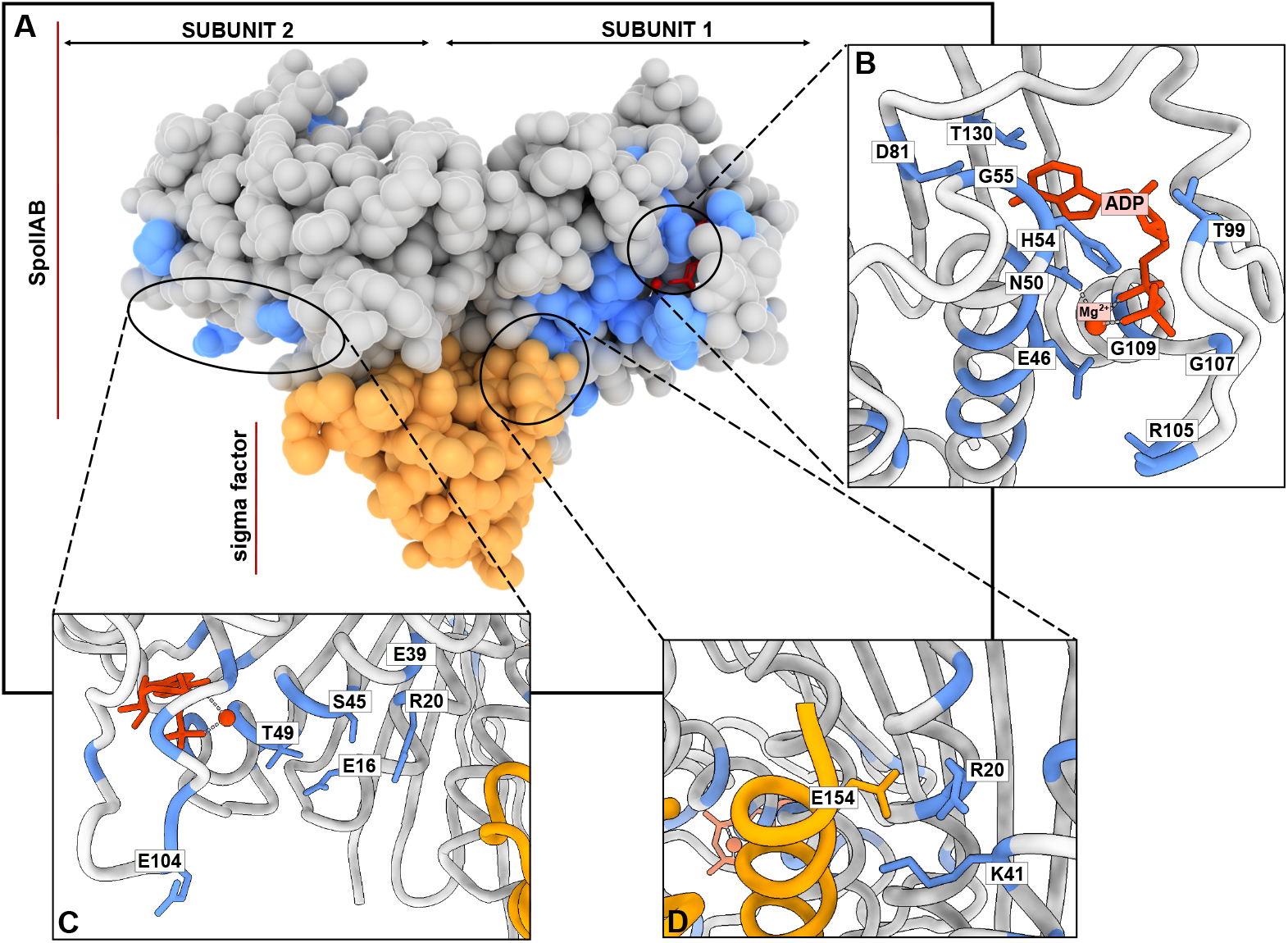
Identifying functional residues in an Anti-sigma F factor (Uniprot O32727, PDB: 1l0O). (A) The dimeric Anti-sigma F factor is shown using Van der Walls surfaces (grey). Predicted functional sites are coloured in blue, Mg^2+^:ADP is shown in red and the bound sigma factor is shown in yellow. (B, C and D) Functional sites identified by our model and previously described in the literature (Campbell et al., 2002; Masuda et al., 2004) are labelled. (B) Predicted functional sites in the proximity to the active site and reported to influence activity. CD) Predicted functional sites involved in interactions with the *σ* factor. (D) Predicted sites that interact with the Anti-sigma F factor antagonist SpoIIAA.

Having shown the model performance using the Anti-sigma F factor as an example, we extended this analysis of functional positions to a larger data set, consisting of 20 enzymes in the Mechanism and Catalytic Site Atlas database (M-CSA, Ribeiro et al. (2018)), as well as five non-enzymes from the Protein-Protein Interaction Affinity Database 2.0 (Vreven et al., 2015). We applied our model to these 25 proteins and identified 16167 SBI variants (15.2% of the total) and assigned 588 residues to be important for function (12.7% of the total positions). We found a small difference in the fraction of predicted functional sites between the enzymes (13.9% of the total positions) and the other five proteins (11.6%). For the 20 proteins in M-CSA, we collected a curated list of 87 residues known to be involved in catalysis (Ribeiro et al., 2018). Given the well-defined functional roles we would expect that most substitutions at these sites would affect the enzymatic function. Our model assigned 62 of these 87 positions (71%) as functional positions (Fig.4A). In 9 out of 20 enzymes the entire set of catalytic residues were classified as functional residues, while in the rest of the dataset the fraction of matching sites ranged between 20% and 80%.

**Figure 4.**
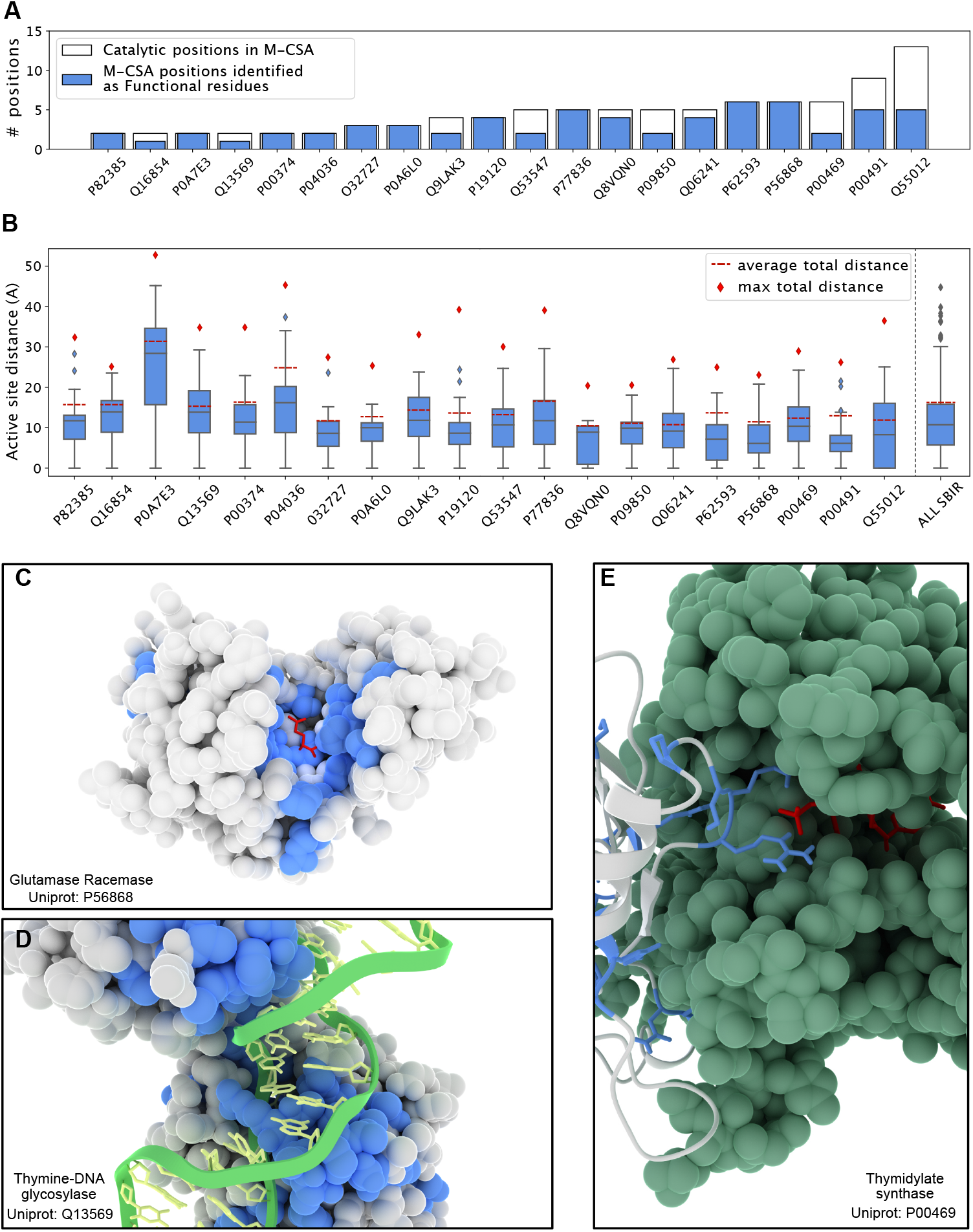
Predicting functional sites in enzymes. We used our model to study functional residues in a set of 20 enzymes from M-CSA, which also provides annotations of residues in catalytic sites. (A) Number of known catalytic residues for each protein in M-CSA (white) and the number of these catalytic residues classified as functional positions by our model (blue). (B) Functional sites are generally close to the active site. For each protein, we show the distribution of distances (using a boxplot) between the predicted functional sites and the (nearest) active site residue (from M-CSA). For comparison we show the average and maximum pairwise residue-residue distances (red dotted line and a red squared dot, respectively). The rightmost boxplot shows the cumulative data. (C, D and E) Examples of predicted functional sites (blue) in three proteins from the M-CSA set. (C) Functional sites in glutamase racemase are found in a single cluster close to the active site. (D) Functional sites in thymine-DNA glycosylase are both located in the active site, but also in the region needed to bind the target DNA chain. (E) Predicted functional sites are also found in protein-protein interfaces, such as in the interface of homo-dimeric thymidylate synthase.

We examined in more detail the 25 of the 87 positions that our model did not assign as SBI. Most of these (17/25 residues; 68%) have a median ΔΔ*G* greater 2 kcal/mol, and our model labelled them as ‘total-loss’ positions. This finding highlights one of the limitations of our approach. While we can assign conserved residues that do not affect stability to have a likely functional role beyond structural stability, the reverse is not true. In these particular cases, ca. 20% (17/85) of the amino acids in the active sites appeared to be important both for function and structural stability. As an example we show the results for an Endo-1, 4-beta-xylanase (Fig.S9) where three of the five catalytic positions were assigned as functional residues by our model, and the remaining two have a median ΔΔ*G >* 2 kcal/mol.

A strength of our model is that it identifies residues with likely functional roles beyond those directly involved in for example catalysis. We examined the positions that our model predicted to be functional sites in the 20 enzymes from M-CSA, and found that a substantial fraction of these are localised in the vicinity of the catalytic site (Fig.4B); on average 48% of the predicted functional residues are located less than 10 Å from the closest catalytic site residue. We expect that many of these residues are important for the catalytic process, as we show for glutamate racemase (Uniprot: P56868, Fig.4C), where 19 out of 26 of the predicted functional residues (including M-CSA catalytic positions) are less than 10 Å from the D-glutamine in the catalytic site (Hwang et al., 1999).

In some cases we also found that the predicted functional sites were located both close to as well as further away (20 Å or more) from the catalytic site, as illustrated by orotate phosphoribosyltransferase (OPRTase, Uniprot: P0A7E3, Fig.S10), where predicted functional residues are located in several distinct regions. Most proteins in the M-CSA set have clusters of predicted functional sites further away from catalytic sites. We found that these positions, for example, may be involved in interactions with substrates, but not directly involved in the catalytic activity. As an example we show a thymine-DNA glycosylase (Uniprot Q13569, Fig4D), where we identified a subset of functional residues far (14.5 Å) from the enzymatic active site. These include Arg275 and the residues in loop 274–277 (Maiti et al., 2009; Fu et al., 2019). These amino acids are not directly involved in catalysis, but their role is to push the target DNA base into the catalytic pocket, which consists of two residues: Arg140 and His151 (Maiti et al., 2009; Kanaan et al., 2015). Our model identified the former as a functional position, while the latter was predicted to be a total-loss position.

In addition to finding positions that are important for catalysis and binding of substrates, our model can also help identify positions that play a role in forming protein-protein interactions as we show for thymidylate synthase (P00469, Fig.4E), a homo-dimeric enzyme with a catalytic pocket in both of the subunits. Of the 41 functional residues identified by our model, 20 are located close to the protein-protein interface. We note here that, unless otherwise noted, all the stability calculations are performed on monomeric structures, and so we identify these residues as being conserved, but not due to the structural stability of the individual subunits. Looking at the predicted functional sites at the interface, we find that some are involved in forming the active site which includes residues from both subunits (Arg178,Arg179,Cys198, Pookanjanatavip et al. (1992)). We, however, also found a number of other residues, which we suggest are conserved as they stabilize the (obligatory) dimer structure.

To examine whether we could extract more information on interaction interfaces we also calculated ΔΔ*G* values using the structure of the dimer of thymidylate synthase, introducing each amino acid substitution in both chains. Together with contact numbers calculated using the dimer structure we use these new scores as input to our model. We compared the resulting classification with the results obtained when using using the monomer structure as input to our model (Fig.5). The analysis shows that nine residues at the interface are classified as total-loss when the calculations are based on the dimer structure (compared to just one when the monomer structure is used). Of these nine total-loss positions, three were labelled as functional residues in calculations based on the monomer structures. Residues at the interface that are known to be important for the catalytic activity (Arg178,Arg179, Pookanjanatavip et al. (1992)) remained labelled as functional sites. We performed a similar set of calculations comparing the classification based on structures of the monomer or dimer structure for OPRTase (Fig.S11A). As for thymidylate synthase, we found differences including an increased number of total-loss positions at the interface in the classification made using the dimer; also in this case the model correctly identified the catalytic residues located at the protein-protein interface.

**Figure 5.**
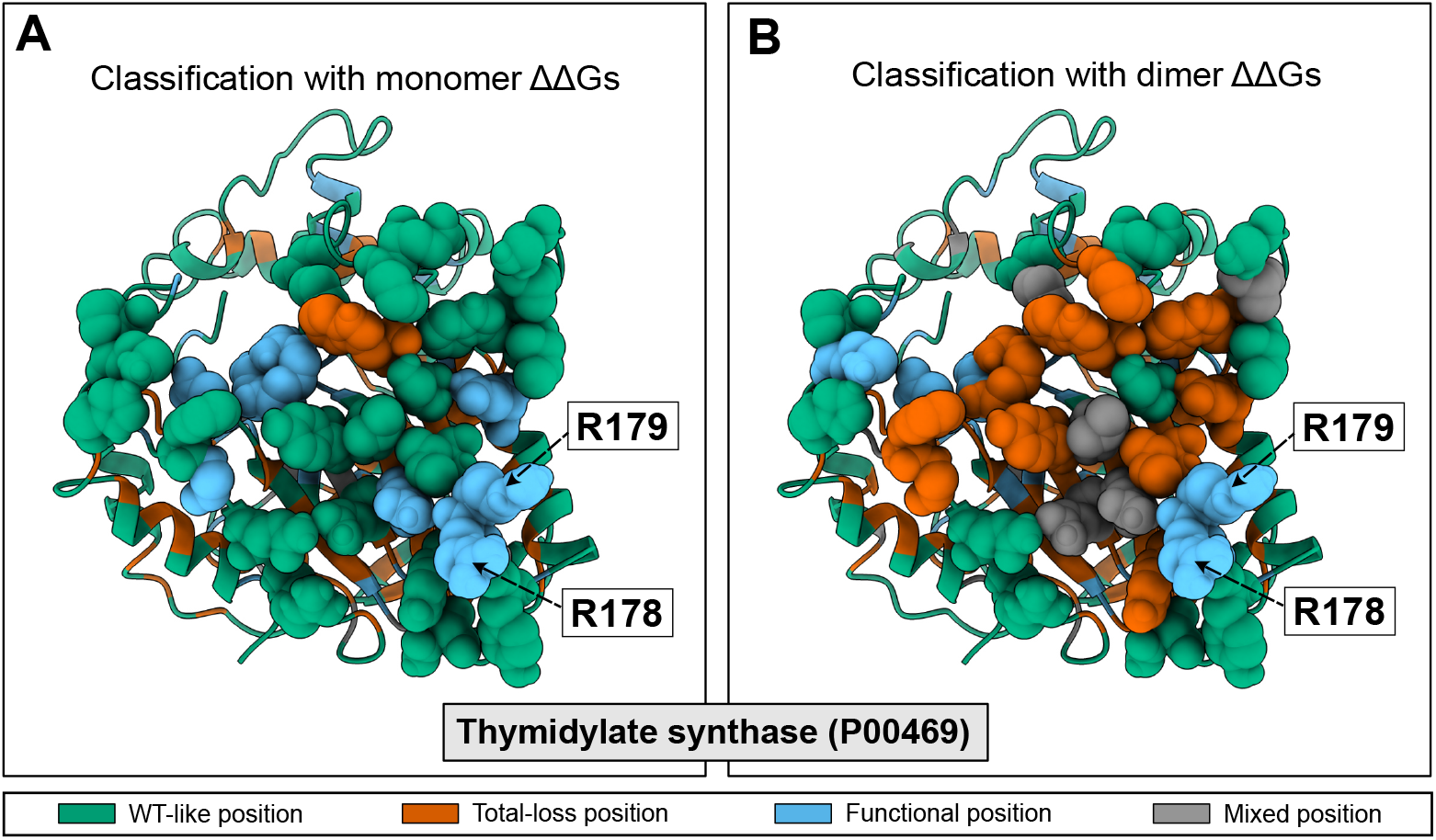
Analysing functional sites at a protein-protein interface. We show the results from predicting functional sites in thymidylate synthase using either the structure of (A) the monomer or (B) the dimer as input to Rosetta for calculating ΔΔ*G*. Residues at the interface are shown with Van der Walls atomic representation. Residues with known catalytic activity are labelled (Pookanjanatavip et al., 1992).

**Figure 6.**
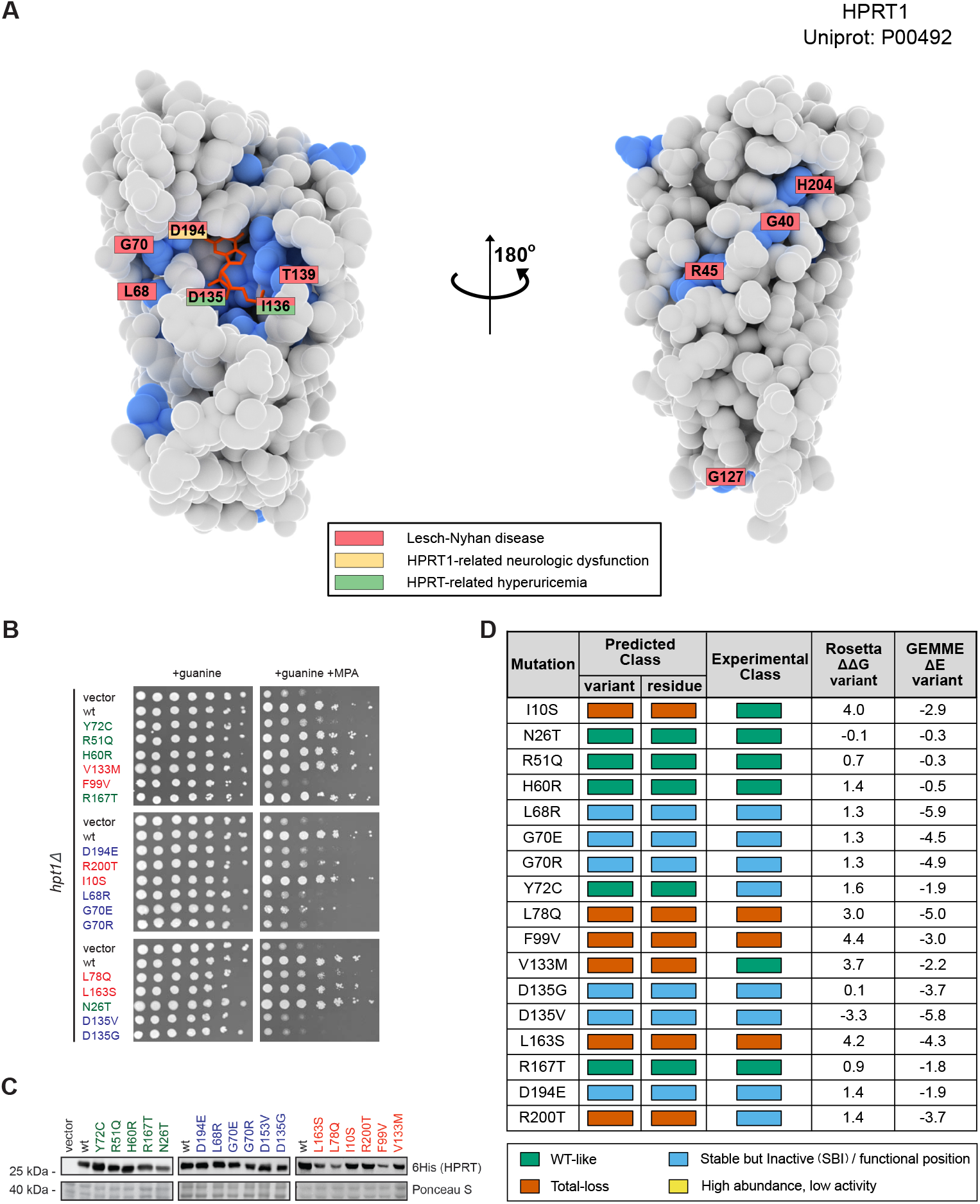
Predicting consequences of missense variants in HPRT1. (A) Structure of HPRT1 shown as van der Walls surfaces with the predicted functional residues coloured in blue. Residues which contain at least one disease-causing variant are labelled and coloured according to which diseases it has been associated with (see legend). (B) Results of a yeast-based assay for HPRT1 function. Yeast strains carrying a vector control, wild-type HPRT1 or one of the 17 HPRT1 variants all grow on medium containing guanine. In the presence of the inosine monophosphate dehydrogenase inhibitor mycophenolic acid (MPA), yeast cells cannot grow in the absence of a functional HPRT1 protein. (C) Assessment of protein abundance using western blots of the 6His-tagged HPRT1 variants. (D) Comparing predictions of the effects of the variants with the experimental measurements. Rosetta ΔΔ*G* values are in units of kcal/mol.

The results described above show that comparing the predictions based on structures of monomers and oligomers can help disentangle functional residues that are key for protein-protein interactions from those with other roles. Specifically, when the structure of the oligomer is known and used as input, our model focuses on residues that are important for functions other than proteinprotein interactions. To illustrate this point further, we compared the classification for (monomeric) myoglobin with that for the *α*_2_*β*_2_ tetramer hemoglobin. For hemoglobin, we made predictions for both the *α* and *β* subunits, and using both the monomer and tetramer structures as input. We first compared the classification results for all three proteins when calculations are based on the monomer structure (Fig.S11C). For all three proteins we found a comparable classification of the residues surrounding the heme group, with 7–9 residues classified as functional. We, however, found differences between myoglobin and the two hemoglobin chains at the residues that form interfaces in the hemoglobin tetramer (Fig.S11C). For these residues, the model assigned the WT-like label for most of the positions in myoglobin, while several interface residues in hemoglobin are classified as functional. This difference likely arises because the calculated ΔΔ*G* values at the interface residues are small (because we used the monomer structures as input), but the residues in hemoglobin are more conserved than those in myoglobin. In line with this hypothesis, many of the interface residues in hemoglobin are classified as total-loss when we use the tetramer structure as input to our model (Fig.S11C). In this case, substitutions at the interface are more destabilizing, and so the model assigns these residues to the class that corresponds to destabilization of the functional protein (the tetramer). Together, the results on thymidylate synthase (Fig.5), OPRTase (Fig.S11AB), and myoglobin/hemoglobin (Fig.S11C) show that residues at key protein-protein interaction sites behave differently depending on whether one considers destabilization of the monomer or oligomer form. In cases where the oligomer structure is known, this difference can help to distinguish residues that play functional roles due to protein-protein interactions from those that are for example directly involved in catalysis. Similarly, when using the monomer structure as input, the model can help shed light on key interaction interfaces, which appear as surface exposed patches of conserved residues.

### Predicting and understanding disease variants in HPRT1

Having validated and exemplified our model using previously published data, we then used our model in a prospective study to predict the impact and mechanism of human missense variants. We and others have previously shown that many, but not all, disease-causing missense variants cause loss of function by loss of abundance. The ability to assign a functional status to so-called variants of uncertain significance is one of the major outstanding challenges in clinical genetics. In order to understand the molecular origin of disease and develop potential treatments it is, however, also important to be able to predict why variants cause loss of function.

We therefore selected hypoxanthine-guanine phosphoribosyltransferase-1 (HPRT1, Uniprot: P00492), an enzyme involved in the onset of Lesch-Nyhan disease and its attenuated variants (Fu and Jinnah, 2012; Fu et al., 2014), for a prospective study of the accuracy and utility of our model. We estimated structural and sequence features for 190 of 210 residues in HPRT1; of the 3610 total possible single amino acid variants, our model predicts 471 to be SBI and 1046 to be total-loss variants.

We then selected 17 variants for experimental characterization. These variants were selected either from gnomAD (Karczewski et al., 2020), ClinVar (Landrum et al., 2018) or variants that we selected to test our model more broadly. Five of the variants were predicted to have wild-type-like activity, six variants were predicted to be total-loss (i.e. loss of activity and loss of abundance), and six variants were predicted to be SBI (i.e. loss of activity without loss of abundance) (Fig6).

To test our predictions, we established a yeast-based growth assay for HPRT1 function. HPRT1 catalyses the formation of inosine and guanosine monophosphate (IMP and GMP) from hypoxanthine and guanine, respectively. Previous studies have shown that mycophenolic acid (MPA) acts as an inhibitor of IMP dehydrogenase, which is responsible for the conversion of IMP to xanthosine monophosphate (XMP, Woods et al. (1983); Escobar-Henriques and Daignan-Fornier (2001)). Thus, in the presence of MPA, yeast cells can only generate GMP through the salvage pathway, and the yeast HPRT1 orthologue, Hpt1, therefore becomes essential (Fig.S12). We introduced each of the 17 variants as well as a wild-type and vector controls into a yeast strain lacking Hpt1 to test their effects on function and abundance. We found that 11/17 variants grew worse that the wild-type control in the presence of MPA, showing that they cause loss of function (Fig6B). We also measured the abundance of the wild-type and 17 variants using western blots and found that 3/17 variants had substantially reduced levels (Fig6C).

We used the experimental data to classify the variants into wild-type-like, SBI and total-loss and compared the results to the computational predictions (Fig6D). Overall we find that 13/17 (76%) of the variants are predicted correctly including all (6/6) of the SBI residues. This result shows that our model can predict variant effects relatively well, and in particular can help separate loss-of-function variants that lose function due to intrinsic function from those due to structural stability. We note that some of these variants have also been characterized biochemically (Fu and Jinnah, 2012), with overall good agreement between our predictions, the yeast assays and the biochemical experiments.

### Making the model more easily accessible

Evaluating ΔΔ*G* values with Rosetta is relatively slow, making the widespread application of our model less straightforward. We have, however, recently developed a method, called RaSP, for rapid stability predictions (Blaabjerg et al., 2022), which uses a deep-learning representation to approximate Rosetta ΔΔ*G* values orders of magnitude faster than Rosetta. We first tested the results when using ΔΔ*G* values from RaSP as input to the functional model described above (which was trained with ΔΔ*G* values generated by Rosetta). Despite the relatively high correlation between the Rosetta and RaSP ΔΔ*G* values (average Spearman correlation coefficient of 0.78), we found that the performance of the model was lower during the cross-validation when we used ΔΔ*G* generated by RaSP (Fig.S13A). This result suggests that differences in the ΔΔ*G* values generated by RaSP and Rosetta combined with the threshold-based structure of gradient boosting machines can shift the prediction for variants with feature values close to the threshold values used inside the model. We therefore retrained our model using instead ΔΔ*G* values generated by RaSP. We found that the RaSP-based model performs as well as the model trained with Rosetta on our validation sets (Fig.S13B) and therefore decided to test the performance for many of the tasks we used to test our Rosetta-based model on. We found that the RaSP-trained model identifies the same number of functional sites as the Rosetta-trained model (62/87) when we examine the enzymes in the M-CSA set, with 57/62 being the same (Fig.S13C). We also found that the RaSP-based model predicts the effects of the HPRT1 variants as well as the Rosetta-trained model (Fig.S13D).

We make our model available via a notebook that can be run using Google Colaboratory (available via https://github.com/KULL-Centre/_2022_functional-sites-cagiada). The notebook guides the user with a step-by-step procedure to generate input data, using for example the GEMME and RaSP web implementations, and to generate the predictions of functional sites.

## Discussion

There is a long history of studying protein function through analyses of sequence conservation, and conserved residues are generally important for the protein. Because most proteins need to be folded to function it is, however, difficult to disentangle the role of individual amino acids on the stability of the overall fold from more direct roles of individual amino acids for protein function. Clearly, it may not always be possible to separate the two, and we have shown for example that substitutions at some catalytic residues may result in loss of stability. In many cases, however, substitutions at functionally important sites either have a small negative effect on stability, or may even give rise to increased stability (Shoichet et al., 1995; Bloom et al., 2006). Building on these ideas we have here presented a method that finds functionally important sites as those that are conserved, but not immediately due to a role in protein stability.

To construct and train our model we leverage (i) large-scale experimental assays reporting on protein activity and abundance and (ii) computational methods to estimate variant effects on protein stability as well as general functional effects using conservation in multiple sequence alignments. We have validated our model using both detailed biochemical experiments on individual proteins (via M-CSA), as well as larger scale data generated by high-throughput experiments. We also validated the method through prospective predictions of the mechanism of disease variants in HPRT1. Our model is freely available. For users that want to avoid slower, more costly Rosetta calculations, we have also trained a model that uses our deep-learning-based method for stability predictions (Blaabjerg et al., 2022). Since both Rosetta (Akdel et al., 2022) and RaSP (Blaabjerg et al., 2022) give relatively accurate results using structures predicted using AlphaFold, our model can also be applied to predicted protein structures. Very recently, Tsuboyama et al. (2022) showed that one can also use large-scale measurements of stability changes as input to an approach inspired by our work.

Our model for functionally important sites, in some sense, lies between two extremes that have previously been used to analyse proteins. At one end of the scale, activity-based MAVEs and sequence conservation analyses provide a global view of residues that play functional roles, but often do not give direct mechanistic insights. At the other end of the scale, detailed biochemical, biophysical and structural experiments are often needed to pinpoint how individual amino acids affect function, but are (with exceptions such as work by Markin et al. (2021)) difficult to scale to a large number of variants and proteins. To test our model we have thus both compared it to data generated by MAVEs, reducing to a one-dimensional view when only a single assay is available, as well as results from detailed assessments of the role of individual amino acids in enzymes and protein-protein interactions.

Technological and methodological developments have led to rapid increases in the number of known protein sequences, as well as our ability to assign overall biological functions to many of these. The ability to identify functionally important sites in proteins is important for our understanding of how proteins work, our ability to engineer enzymes and to understand the mechanisms that underlie diseases. Our model may, for example, be used to select residues for protein engineering experiments, or for generating focused libraries in enzyme optimization. As additional data is generated by MAVEs reporting on both function and abundance, the model can be improved further. As we have illustrated, the model can help pinpoint the molecular mechanisms of disease variants, which may both guide further experiments and be used as starting point for developing mechanism-based therapies such as finding variant that may or may not be rescued by pharmacological chaperones. Recent work shows how it is possible to annotate enzyme function at larger scale (Yu et al., 2023) and our model can help pinpoint functionally important sites in newly discovered enzymes. We thus anticipate that the method that we have described here will help researchers make progress in these and many other areas.

## Methods

### Preprocessing of the training data

The training data include paired MAVE data for three different proteins: NUDT15, PTEN and CYP2C9. For each protein, we selected thresholds for the scores generated by each MAVE as outlined previously (Cagiada et al., 2021). Briefly, we fit the variant score distributions to three Gaussians and use the intersection of the first and last Gaussian as the cutoff. We use such binary classification for each of the two MAVEs to classify the variants into four categories (Cagiada et al., 2021).

### Sequence features

For each protein in the training (TableS1) and validation (TablesS4,S5andS6) data we performed a statistical analysis of multiple sequence alignments (MSA). We extracted the sequence of the first isoform from Uniprot and used HHBlits (Remmert et al., 2012) to build an MSA, on which we applied additional filters before evaluating the substitution effects and conservation scores. The first filter keeps only the positions (columns) that are present in the target sequence, and the second filter removes the sequences (rows) where the number of total gaps exceeds 50% of the total number of positions. We used GEMME (Laine et al., 2019) to calculate an evolutionary conservation score. GEMME explicitly models the evolutionary history of the protein, returning a score (ΔΔ*E*) which estimates an ‘evolutionary distance’ of a variant from the query wild-type sequence. The ΔΔ*E* scores for the variants range between 0 (conservative substitution) and −7 (substitutions that appear incompatible based on the MSA). In addition to the effect of the individual variants, we also calculated a ‘neighbour score’ as the average of the ΔΔ*E* values for the previous and proceeding residue. In addition to the ΔΔ*E* scores, we used the hydrophobicity (Monera et al., 1995) of the target amino acid as a feature in our model.

### Structural features

We predicted changes in thermodynamic stability using Rosetta (GitHub SHA1 99d33ec59ce9fcecc5e4f3800c778a54afdf8504). We used the Cartesian ddG protocol (Park et al., 2016) on crystal structures corresponding to the Uniprot sequence listed in TablesS1,S4 andS5. The ΔΔ*G* values obtained from Rosetta were divided by 2.9 to bring them from Rosetta energy units onto a scale corresponding to kcal/mol (Park et al., 2016). Stability changes using RaSP were performed as described (Blaabjerg et al., 2022). We used these ΔΔ*G* values as a feature for the classifier, setting all values below/above a range of 0 – 5 kcal/mol, to 0 or 5 kcal/mol, respectively. We also calculated a neighbour score, averaging over the two nearest neighbours, as for the ΔΔ*E* values.

Finally, we used MDTraj v.1.9.3 (McGibbon et al., 2015) to calculate the weighted contact number (WCN) for each residue:

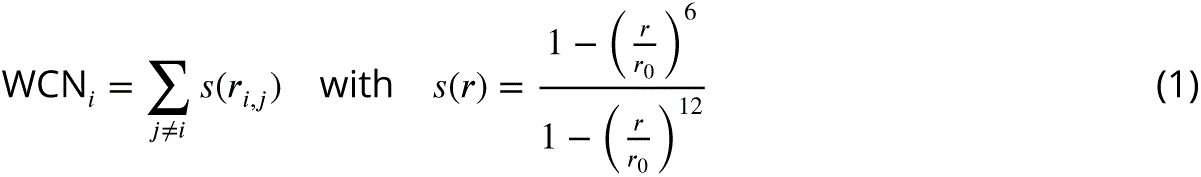

where *r_i,j_* is the C*_α_*distance between residue *i* and *j*, *r*_0_ is a switching parameter (set to 7.0 Å).

We determined solvent exposed and buried regions using GetArea (Fraczkiewicz and Braun, 1998) using 20% as a threshold to divide the data in to two classes.

### Training the classifier

We began by including up to eleven features as input to our classifier. After testing on the three proteins (NUDT15,PTEN and CYP2C9), we proceeded with only eight of them. The features used for each variant were: (1, 2) variant ΔΔ*G* and ΔΔ*E*, (3–6) residue average and neighbour average ΔΔ*G* and ΔΔ*E* scores, (7) the hydrophobicity of the target amino acid, and (8) the WCN. The three features discarded during model building were: wild-type amino acid type, binary exposure classes based on solvent exposure and hydrophobicity of the wild-type amino acid.

We used a gradient descent machine to classify the variants, as implemented in Catboost (Prokhorenkova et al., 2017) v.0.26.1, using a Multinomial/Multiclass Cross Entropy Loss and L2 regularization. We first set the number of gradient descent iterations (Fig.S2A) on the ‘vanilla’ model and then used a grid scan to find optimal value for the remaining hyperparameters. We measured precision and recall using five-fold cross validation on the training data to optimise the hyperparameters (Fig.S2B). We compared the model performance with a null model from the sklearn python package (*sklearn.dummy.DummyClassifier*) using the ‘prior’ strategy, which returns the most frequent class label in the observed argument passed to it. We also used a random forest classifier model (*sklearn.ensemble.RandomForestClassifier*) in the comparison, optimizing the hyperparameters using a grid scan protocol and *k*-fold cross validation.

### Cloning

Full-length human HPRT1 was expressed in yeast from the pYES2 vector (Invitrogen). An N-terminal RGS6His-tag was inserted upstream of an SRS linker peptide, before HPRT Met1. Point mutations were generated by Genscript.

### Yeast strains and techniques

The *hpt1*Δ (MatA, *his3*Δ*1, leu2*Δ*0, met15*Δ*0, ura3*Δ*0, hpt1::KanMX*) *Saccharomyces cerevisiae* yeast strain was from Invitrogen. The cells were cultured in synthetic complete (SC) medium (2% glucose, 6.7 g/L yeast nitrogen base without amino acids and with ammonium sulfate (Sigma)) and supplemented for selection with 1.92 g/L uracil drop-out mix (US Biological). For expression, the glucose was replaced with 2% galactose.

Yeast transformations were performed with lithium acetate (Gietz and Schiestl, 2007). For growth assays, the cells were cultured at 30 ^◦^C to exponential phase and diluted to an OD(600nm) of 0.40. From this, dilution series (5-fold) were prepared and 5 *µ*L of each dilution was applied as droplets on agar plates. Colonies formed after 2–3 days of incubation at 30 ^◦^C. The agar contained 50 *µ*g/mL guanine (Sigma) and 0.1 mg/mL MPA (Sigma).

Whole cell lysates for western blotting were prepared from exponential phase cultures using glass beads and trichloroacetic acid (TCA) as described before (Kampmeyer et al., 2022). Briefly, · 10^8^ cells in exponential phase were harvested and washed in water by centrifugation (3000 g, 5 min.). The cells were resuspended in 1 mL of 20% TCA and centrifuged (3000 g, 5 min.). The supernatant was discarded and the pellet was resuspended in 200 *µ*L 20% TCA and transferred to 2 mL screwcap tubes containing 0.5 mL 400-600 micron glass beads (Sigma). The tubes were then applied to a Mini-BeadBeater machine (BioSpec Products Inc.) set at 3 cycles of 10 s. With a needle, a hole was made in the bottom of the tube and the tube was placed inside a 15 mL tube. Then, 400 *µ*L 5% TCA was added and the material was eluted by centrifugation (1000 g, 5 min.) into the 15 mL tube. The eluted material was centrifuged (10000 g, 5 min.) at 4 ^◦^C. The pellet washed twice with ice-cold 80% acetone. Finally, the pellet was resuspended in 100 *µ*L sample buffer for SDS-PAGE (62.5 mM Tris/HCl pH 6.8, 2% SDS, 25% glycerol, 0.01% bromphenol blue, 5% *β*-mercaptoethanol) and incubated for 5 min. at 100 ^◦^C.

### SDS-PAGE and western blotting

SDS-PAGE was performed using 12.5% acrylamide gels. After electrophoresis, the proteins were transferred to 0.2 *µ*m nitrocellulose membranes (Advantec) by electro-blotting. After transfer, the blots were stained with Ponceau S (0.1% Ponceau S in 5% acetic acid) and blocked in PBS (10 mM Na_2_HPO_4_, 1.8 mM KH_2_PO_4_, 137 mM NaCl, 3 mM KCl, pH 7.4) with 5% skimmed milk powder. The antibodies were anti-RGSHis (Qiagen, Cat. No. 34650) and peroxidase-conjugated anti-mouse antibody (Dako, Cat. No. P0260).

## Data and code availability

Data available at https://github.com/KULL-Centre/_2022_functional-sites-cagiada.

## Acknowledgments

The research was supported by the PRISM (Protein Interactions and Stability in Medicine and Genomics) centre funded by the Novo Nordisk Foundation (NNF18OC0033950, to A.S., R.H.P. and K.L.-L.). We acknowledge access to computing resources from the Biocomputing Core Facility at the Department of Biology, University of Copenhagen.

## Supplementary Material

### Supplementary Figures

**Figure S1.**
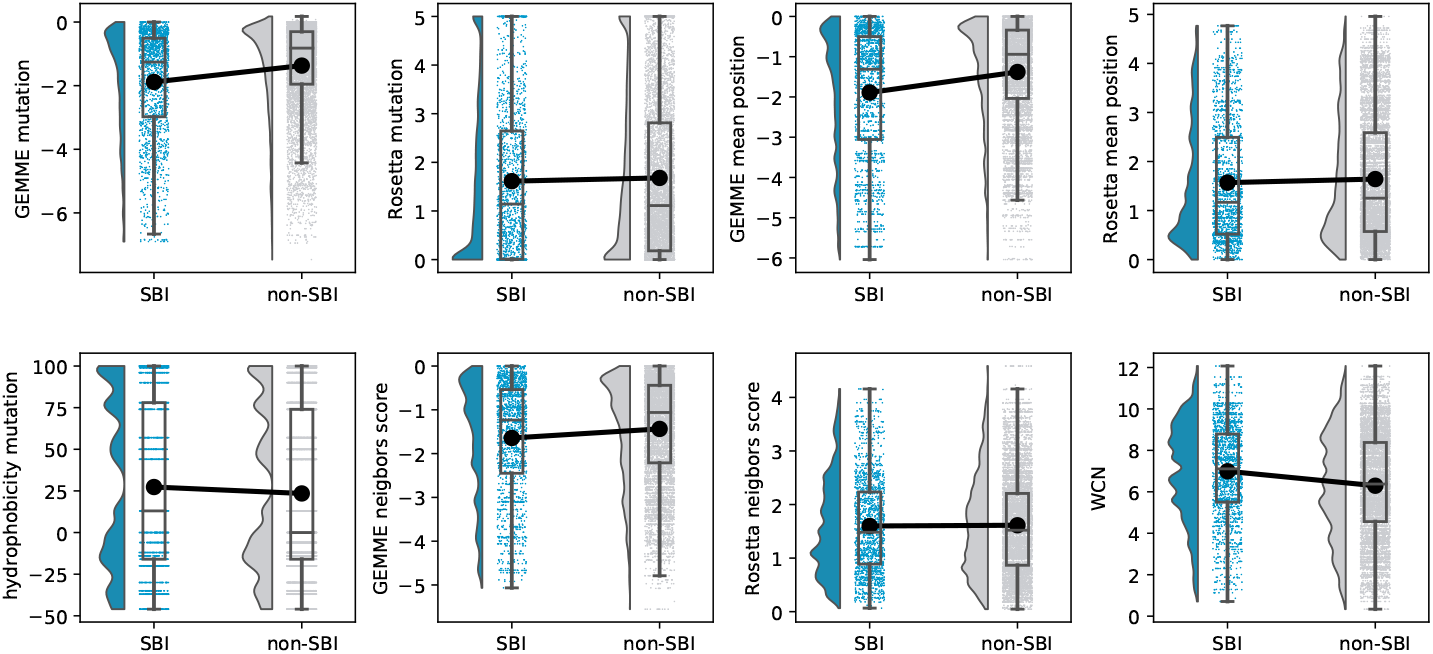
Comparing values of the individual features for SBI and non-SBI variants. Each raincloud plot shows, for each feature used in the model, the distribution of feature values in the training proteins for variants belonging to the SBI class (as assigned by experiments) or in one of the other three classes (WT-like, total loss and ‘low abundance, high activity’). Each plot also shows the median, quartiles with a box plot and comparison between median values with a black line. Raw data are also displayed as points under the box plot.

**Figure S2.**
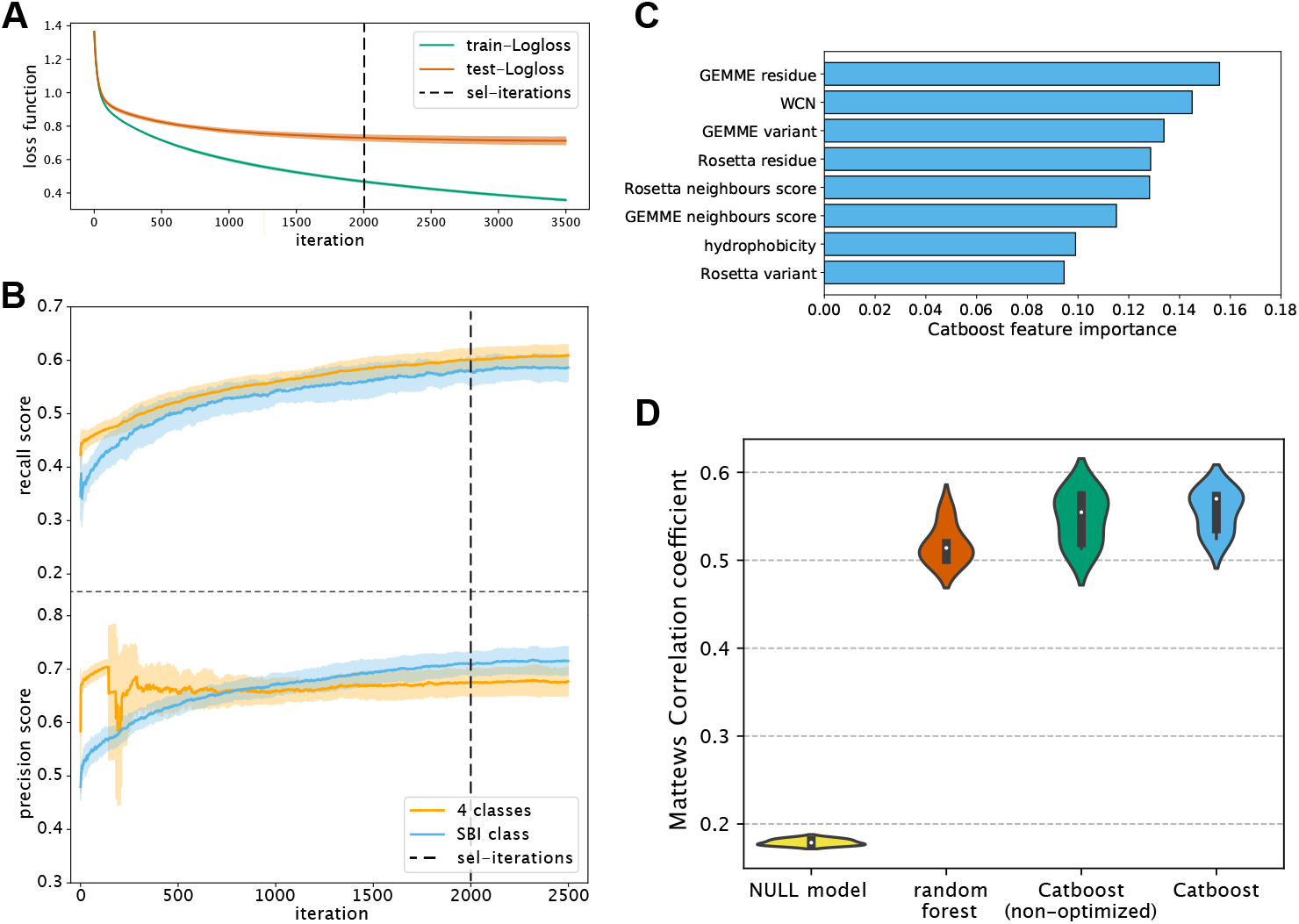
Details of the variant classifier training process. (A) Changes in the loss function for the (green) training data and (red) test data during model training. (B) Progress of performance (measuring recall and precision on the test set) of the Catboost classifier during training. The yellow lines represent the recall and precision for correctly classifing the variant in one of the four classes, while the blue lines show the performance of the model for classifing the SBI variants. The shaded areas in panels A and B represent the standard deviation from the *k*-cross validation procedure on the training dataset, and the selected number of iterations is shown as a black vertical line. (C) Feature importance for each of the features used to train the model. (D) Comparison of the performance of the Catboost model (using the Matthews’ correlation coefficient), a Catboost version before optimizing hyper parameters, an optimized Random Forest model, and a Null model (which always returns the most frequent class label). The white points inside the ‘violin plots’ represent the median values and the black squared areas represent one standard deviation.

**Figure S3.**
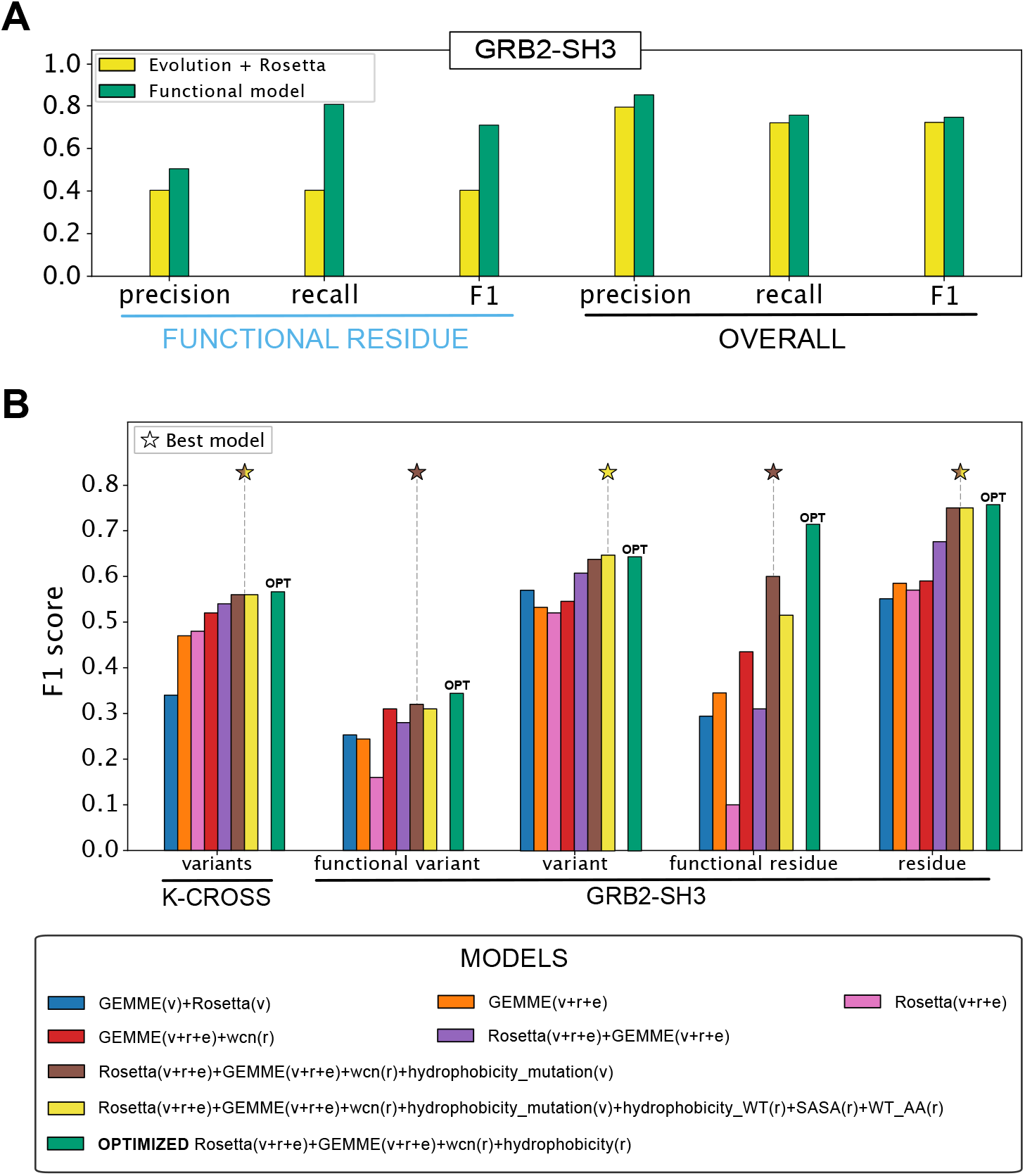
Benchmarks of our functional sites model. (A) Comparison of the predictions of residue classes in the GRB2 SH3 domain using either our optimized model (green) or simply using cutoff values for evolutionary conservation and thermodynamic stability changes (yellow). The three leftmost bars show the results for the subset of functional residues, while the three rightmost series report the results for all the positions predicted. In both the cases precision, recall and F1-score are used as metrics. (B) Comparison between results from our vanilla model (in brown) with vanilla models trained using other sets of features. Results from our final version with optimized hyperparameters are reported (in green). F1 score is shown on the y-axis and the stars highlight the best model for each set (excluding the fully optimized model). The leftmost set reports the results for the cross validation made on the training/test set, while the other sets show the scores for GRB2 SH3 domain, both for the functional residue subset and for all the residues. The legend reports which features were used with v, r and e representing variants, residues and environment, respectively (see Methods for list of features).

**Figure S4.**
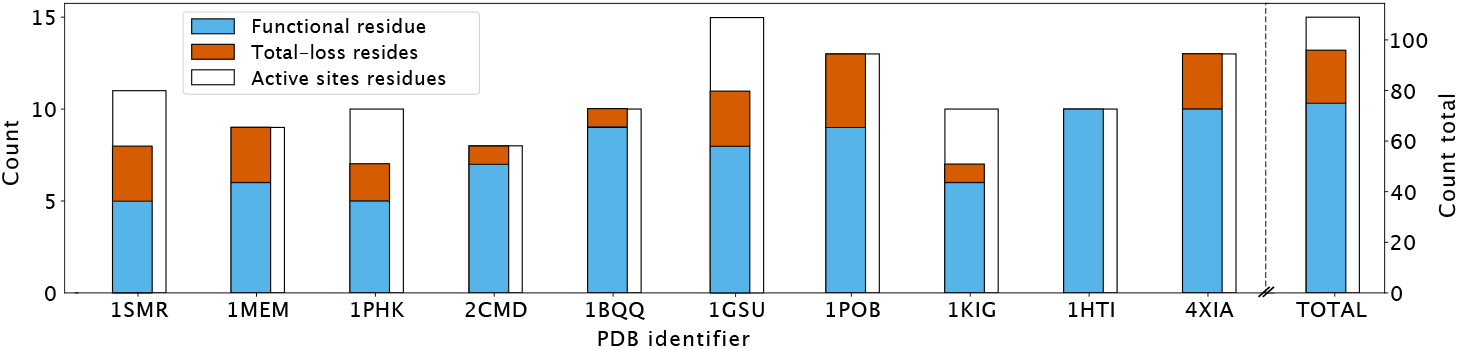
Prediction of active site residues for the set of ten enzymes from Chelliah et al. (2004). The figure shows for each enzyme in the dataset the total number of reported active site positions (white), the subset of these predicted as being functional residues by our model (blue) and the positions predicted to be total loss (red). The rightmost bar shows the cumulative data.

**Figure S5.**
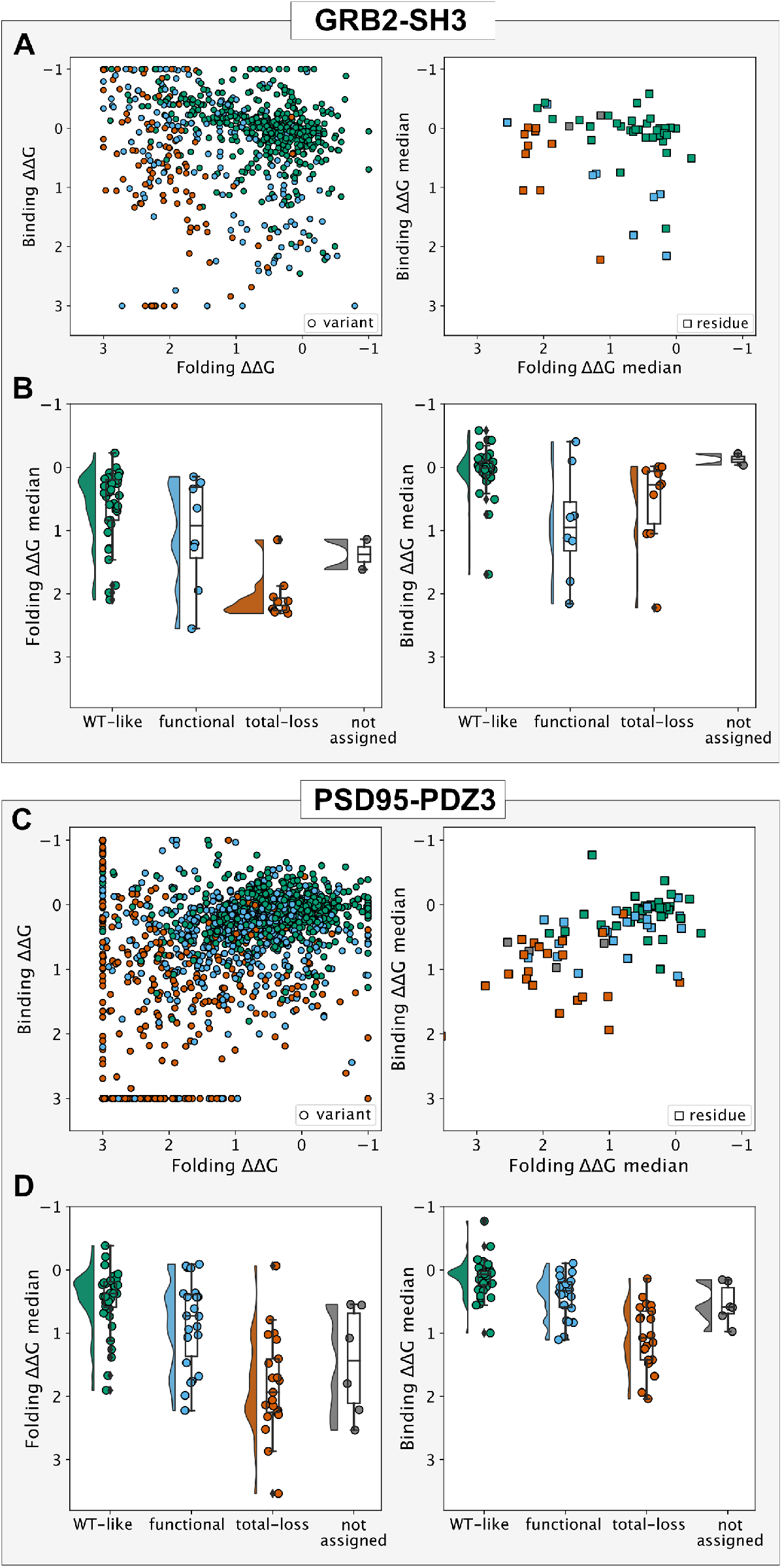
Comparison of experiments and predictions for the GRB2-SH3 and PSD95-PDZ3 domains. (A, C) For each of the two domains we used our model to classify variants and residues and compared to ΔΔ*G* values for domain stability (Folding ΔΔ*G*) and peptide binding (Binding ΔΔ*G*) inferred from experiments. Variants and residues are coloured according to the predicted class (WT-like: green; SBI/Functional site: blue; Total-loss: red; Not assigned: grey). (B, D) Rain-cloud plots of the median value of the folding or stability ΔΔ*G* values for different positions divided into the different classifications. The box plots show the median and quartiles (of the residue median values).

**Figure S6.**
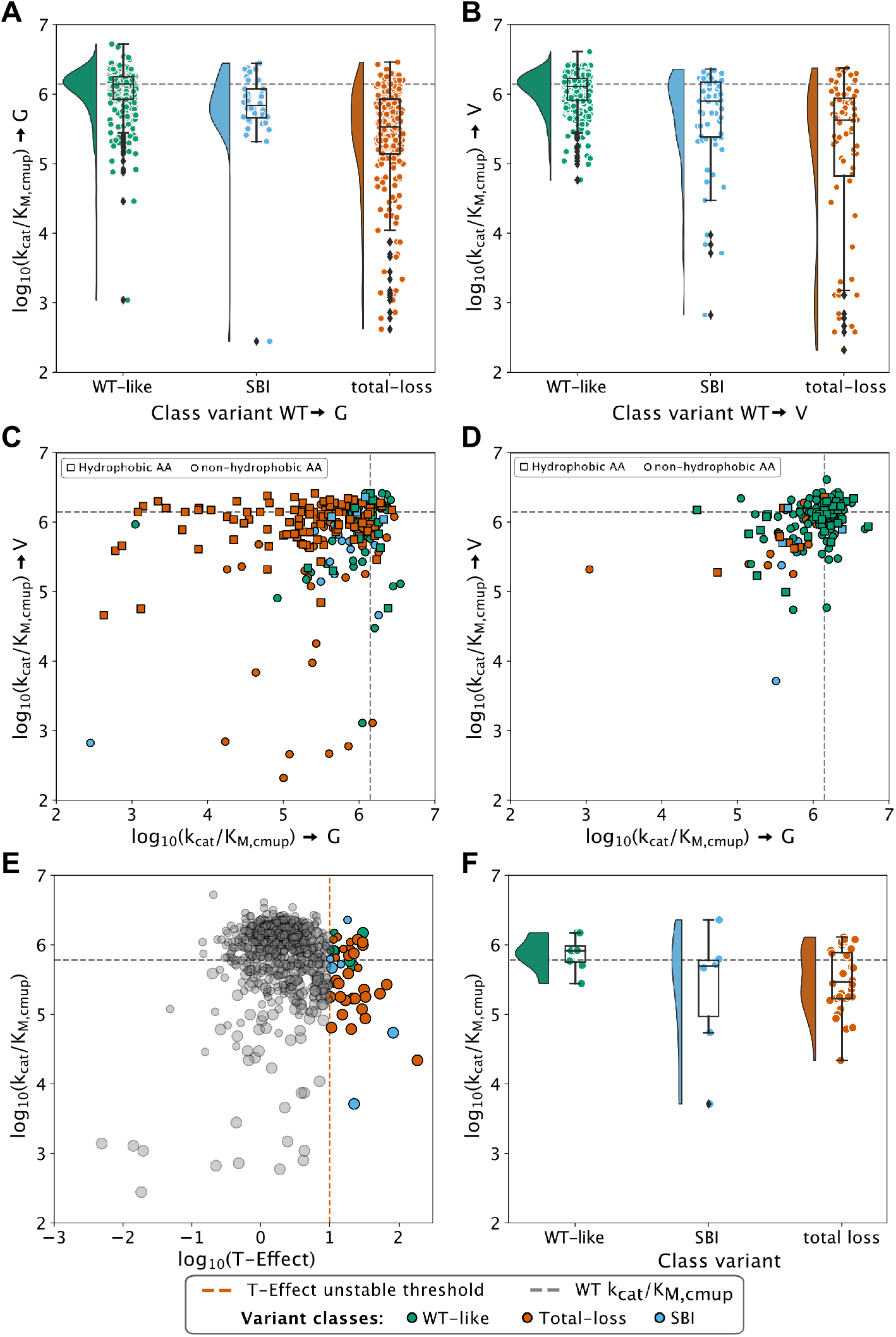
Comparison of experiments and predictions for PafA (Uniprot: Q9KJX5). Raincloud plots for experimentally determined *k*_cat_/*K*_M,cMUP_ values when the wild-type amino acid was was substituted for (A) glycine or (B) valine. In both A and B, the values are shown separately for the different classes predicted by our model. The box plots shows the median and quartiles for each set of variants. The dashed horizontal lines show *k*_cat_/*K*_M,cMUP_ for WT PafA. (C, D) Scatter plot of *k*_cat_/*K*_M,cMUP_ values when substituting a residue with either glycine or valine. (C) Shows the comparison for buried residues (with an exposed surface area of less than 20%) and (D) shows exposed residues. In both C and D squared markers represent positions where the WT residue is hydrophobic and circles indicate non-hydrophobic WT residues. (E) Comparison of experimental *k*_cat_/*K*_M,cMUP_ values and temperature effects during expression. In particular, the T-effect value represents the change in measured catalytic efficiency when the protein was expressed at 23 ^◦^C or 37 ^◦^C (but with the enzymatic assay performed at 23 ^◦^C in both cases). Variants that show a substantial (greater than 10-fold) change nearly all belong to the total-loss category. Points with larger markers represent data for which the T-effect might be underestimated due to experimental limitations. (F) Raincloud plots for the coloured points in panel E (large T-effect).

**Figure S7.**
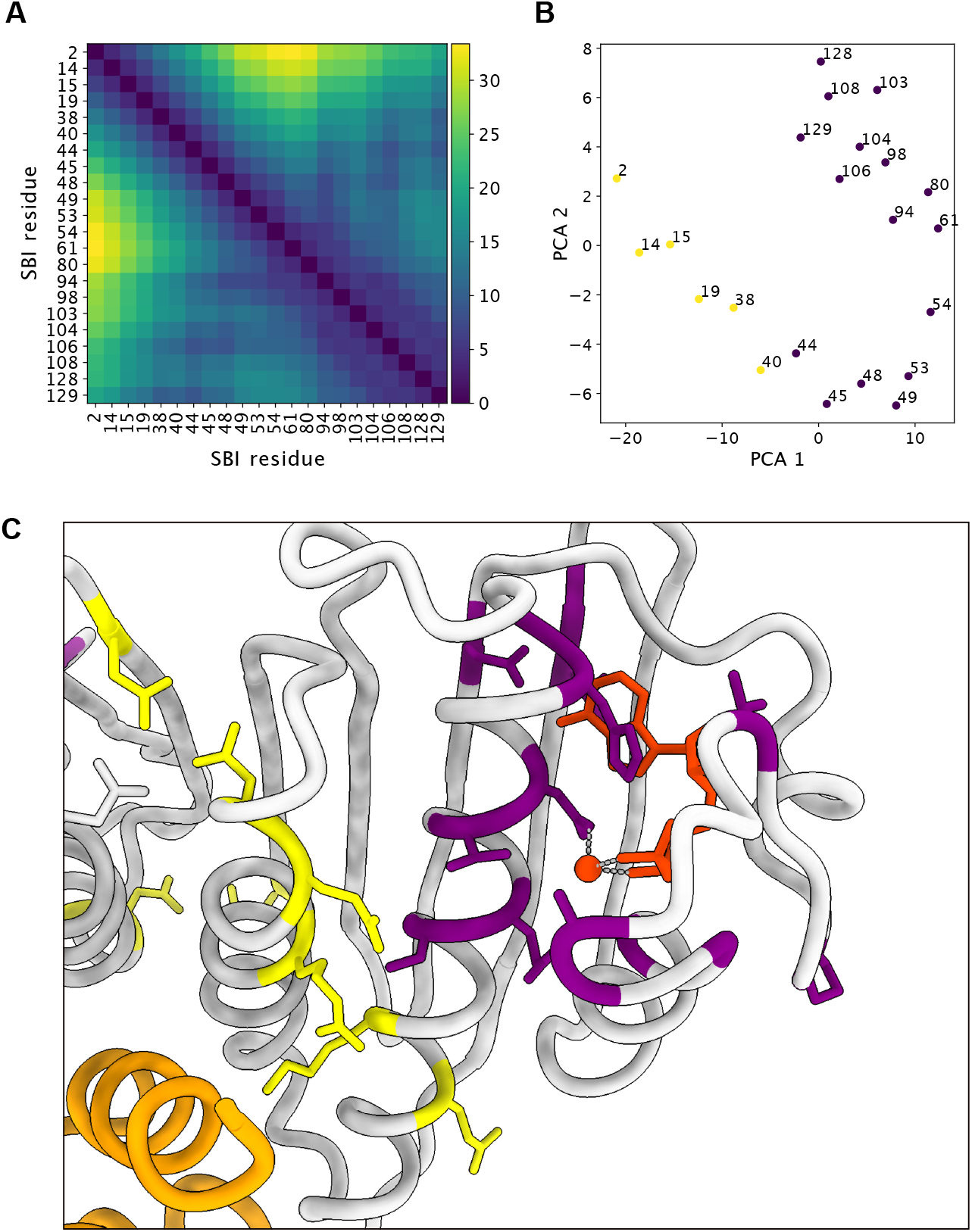
Clusters of functional positions in Anti-Sigma factor. (A) C*_α_* distance matrix for the positions classified as functional by our model. (B) A principal component analysis of the distance matrix suggests that the predicted functional sites can roughly be separated into two spatial clusters (coloured in yellow and purple along the first two principal components). (C) Functional sites coloured using the clustering.

**Figure S8.**
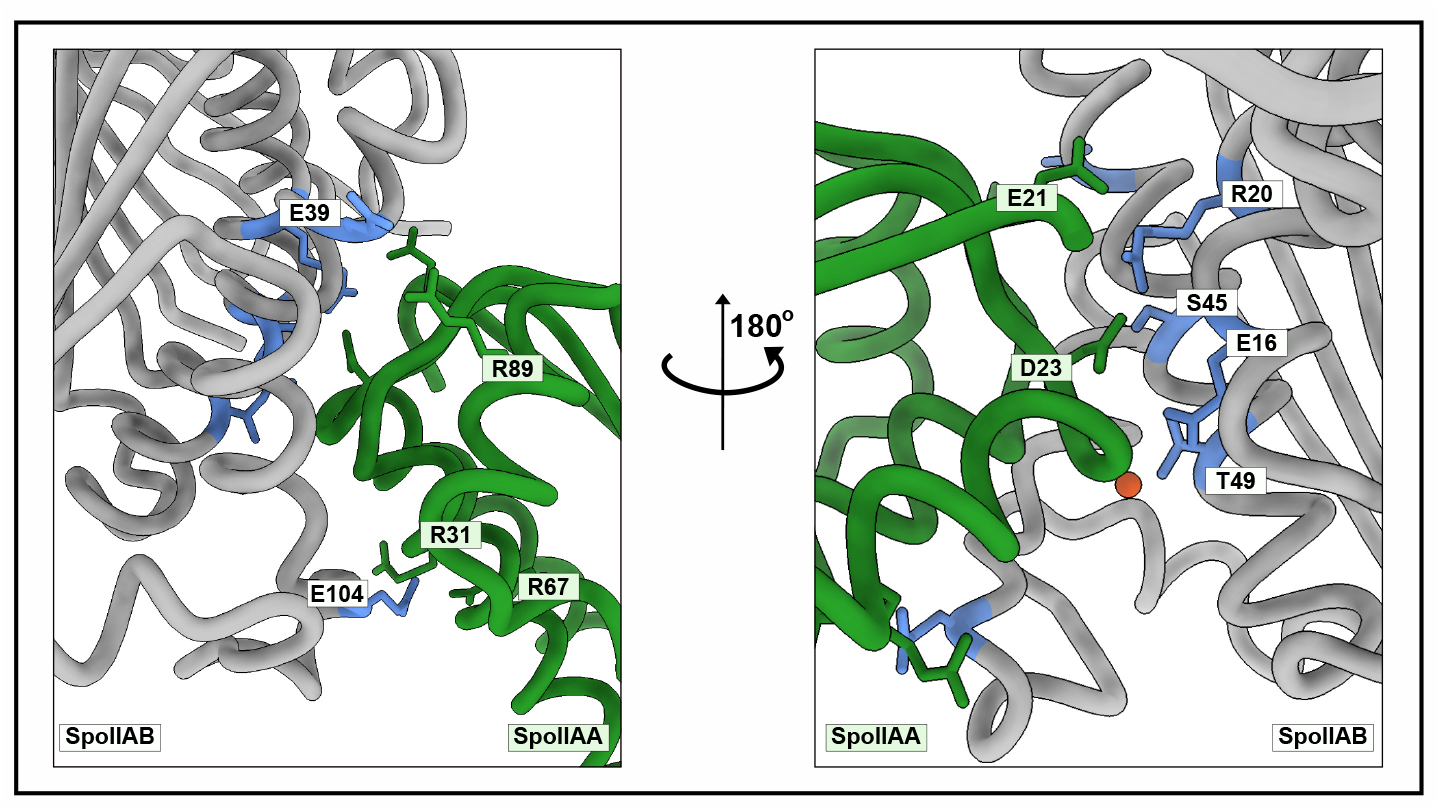
Map of detected functional residues at the interface between the Anti-sigma factor (SpoIIAB in grey) and its antagonist (SpoIIAA in green). Predicted functional residues in SpoIIAB involved in interface interactions are labelled and coloured in blue.

**Figure S9.**
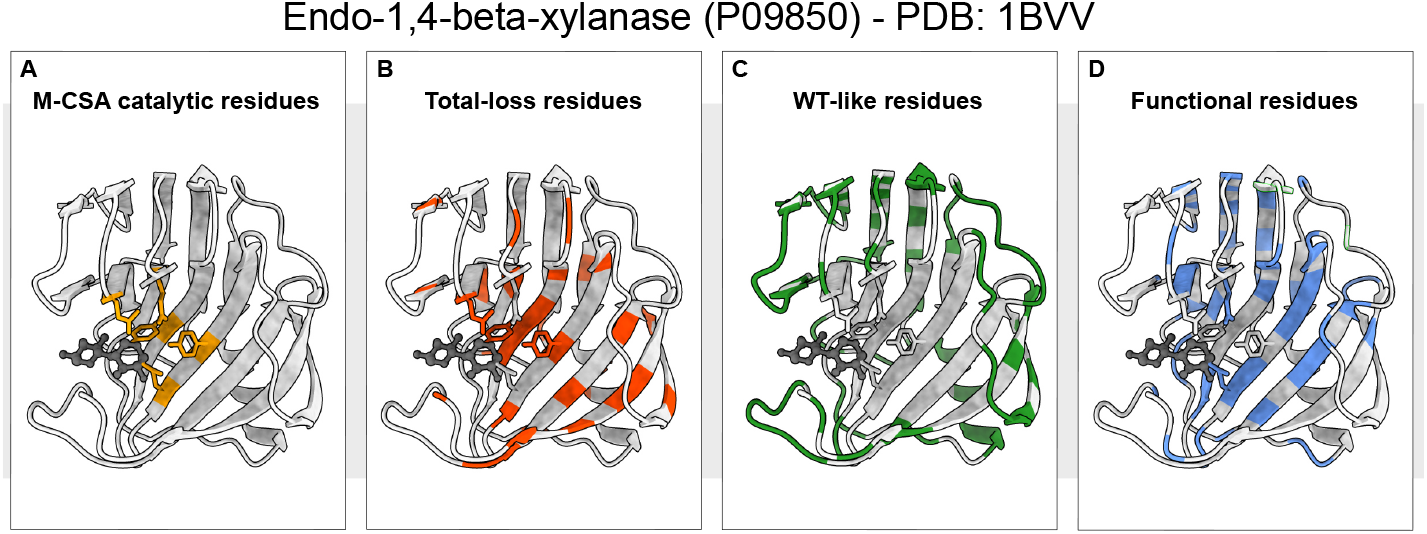
Model predictions for Endo-1, 4-beta-xylanase. (A) Catalytic residues in Endo-1, 4-beta-xylanase as assigned by M-CSA. (B, C and D) show the residues which are classified by our model as (B) total loss, (C) wild-type like, and (D) functional residues. Our model predicts that substitutions at all five catalytic residues affect the protein function rendering it inactive. Three of the five positions are predicted to be important for function, but not for protein stability, whereas two positions are predicted to be important for both function and stability.

**Figure S10.**
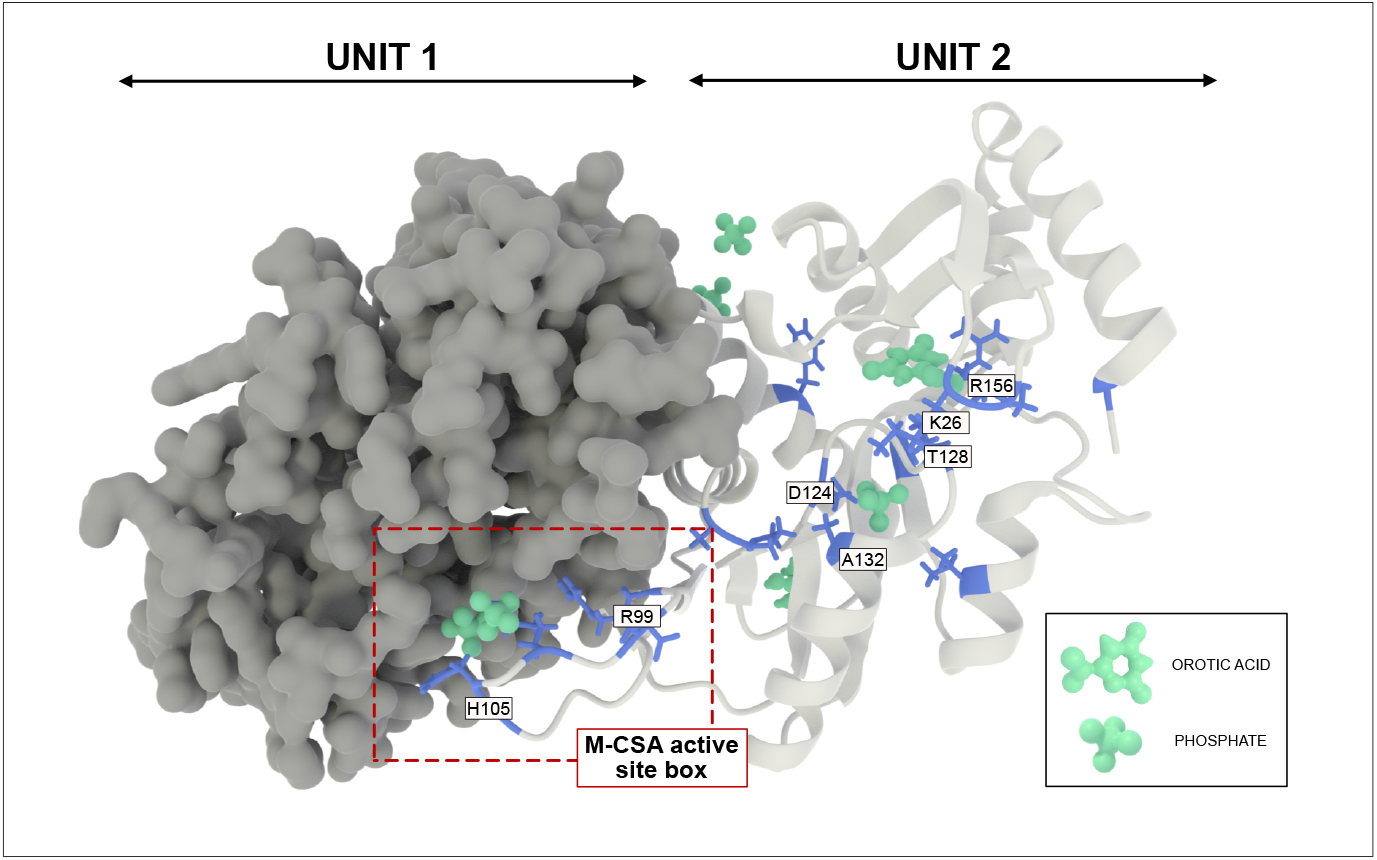
Location of predicted functional sites in OPRTase. The figure shows the dimer structure of OPRTase, with the two sub-units coloured in grey and white, respectively. The residues classified as functional by our model are coloured in blue and labelled on the second sub-unit. The region surrounded by a dotted red box contains catalytic residues reported in the M-CSA database (again shown for the second sub-unit). Orotic acid and phosphate ions bound to the OPRTase complex are coloured in green.

**Figure S11.**
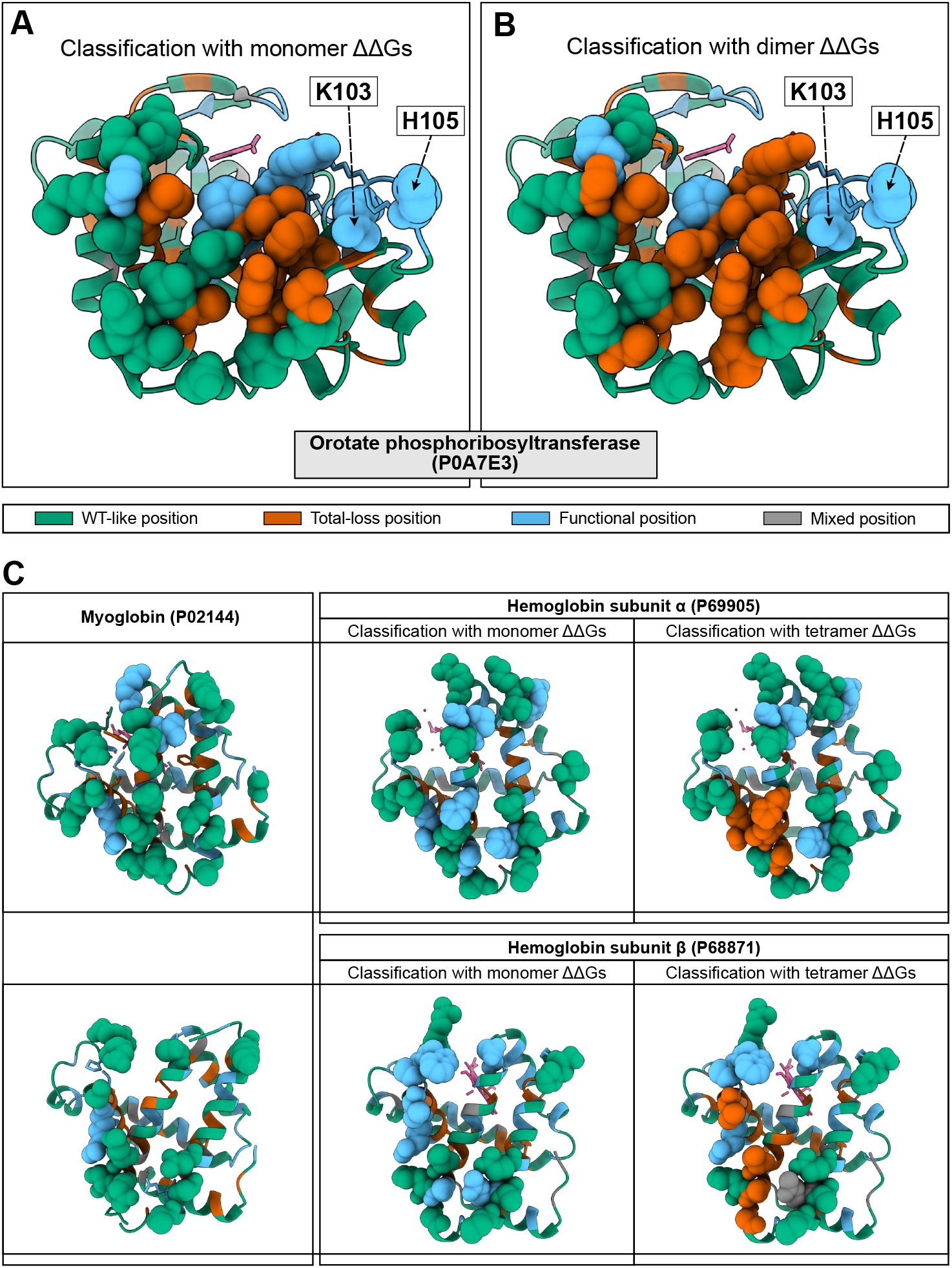
Additional examples of the effect of input structure choice on residue classification in oligomeric proteins. Panels A and B show differences of classification for residues in orotate phosphoribosyltransferase when we use either (A) the monomer or (B) dimer structure as input to the Rosetta ΔΔ*G* calculations. Residues at the interface are shown with van der Walls atomic representation and residues involved in forming the active site at the dimer interface are labelled. Panel C shows, like panel A, a comparison of predictions for human myoglobin and the *α* and *β* subunits of human hemoglobin. For human hemoglobin, the left column shows the residue classification using ΔΔ*G* from the monomer, while the right column the classification made with ΔΔ*G* keeping the entire tetrameric structure during the evaluation. Residues at the tetramer interface are shown with van der Walls atomic representation in all the hemoglobin panels; residues at the corresponding positions in myoglobin are highlighted to make comparisons easier.

**Figure S12.**
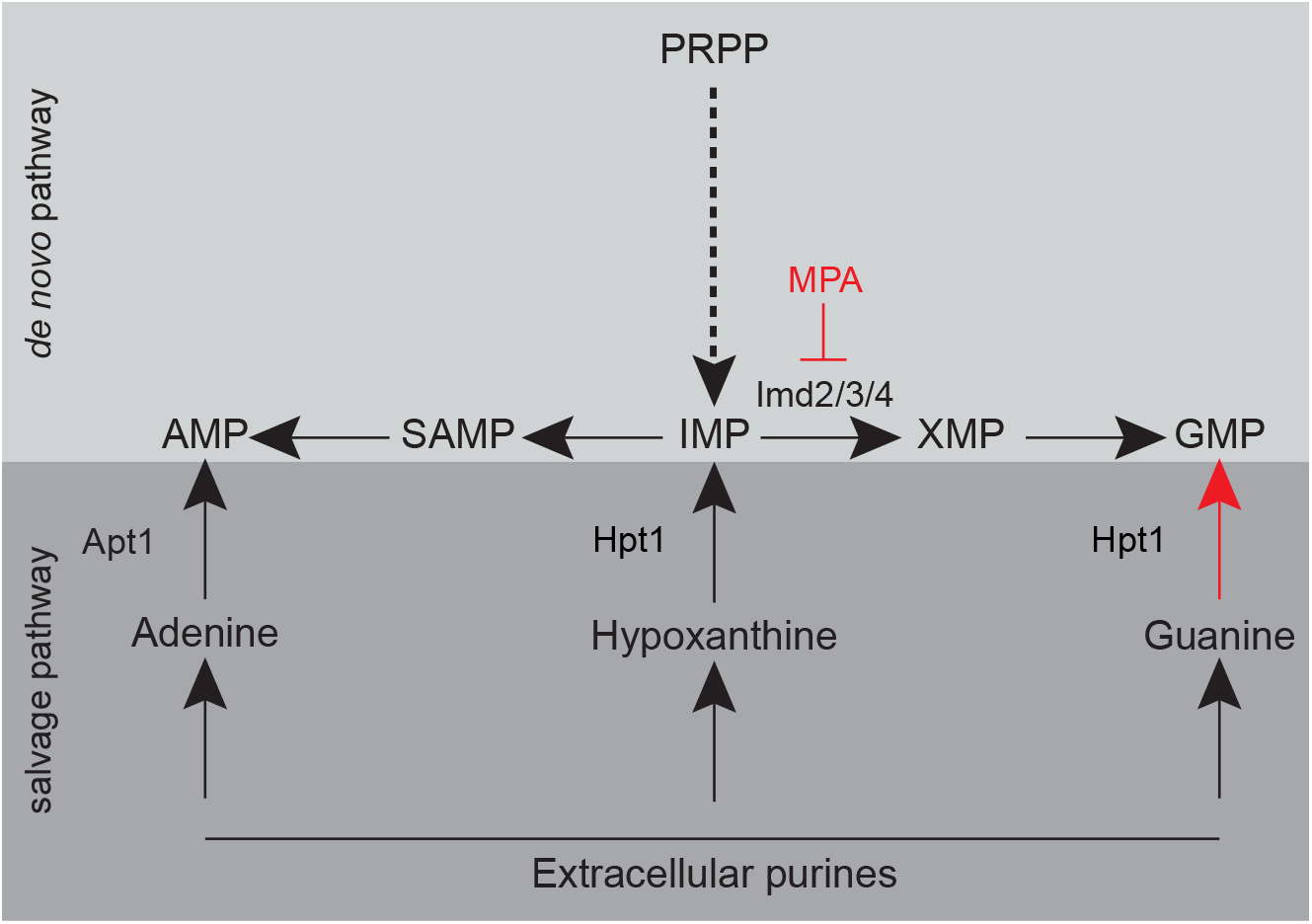
*Hpt1* is essential in the presence of MPA. The figure shows relevant parts of yeast nucleotide metabolism including the salvage pathway where *Hpt1* is essential for generating GMP when XMP synthesis is blocked by MPA. A *hpt1*Δ yeast strain can therefore not grow in the presence of MPA, but can be rescued by human HPRT1 or functional HPRT1 variants.

**Figure S13.**
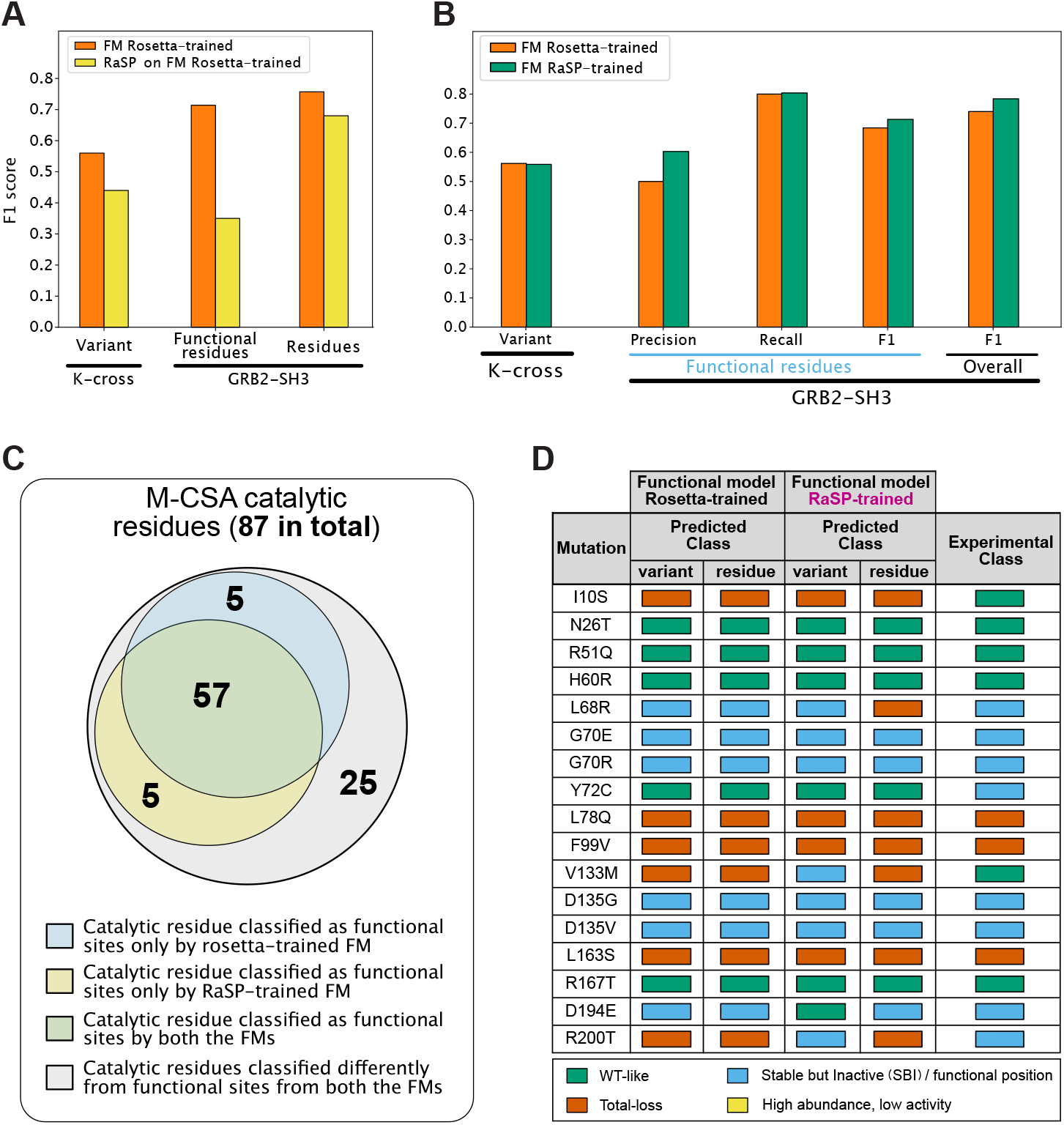
Training a model for functional sites using thermodynamic stability changes predicted by RaSP. (A) Effects of using RaSP ΔΔ*G* predictions as input on the functional residue model trained with Rosetta ΔΔ*G* values. The leftmost set reports the F1 score on the test set during cross validation using as ΔΔ*G* from Rosetta or RaSP. The two rightmost sets show the prediction performances on the GRB2 SH3 domain, for both the functional subset and over all the positions. (B) Comparing the performance of functional residue models trained using ΔΔ*G* data from Rosetta (in orange) and RaSP (in green). The leftmost set reports the F1 score for the cross validation on the training data. The performance on the SH3 domain are shown in the remaining sets. (C) Venn diagram comparing the residues in the M-CSA set predicted from the two models. (D) Predictions of variant and residue classes by the two models are compared to the experimental results on the HPRT1 (rightmost column).

### Supplementary Tables

**Table 1.** Validation using multiplexed assays of variant effects.

| Protein | PDB | MAVE | SBI accuracy | Global accuracy |
| --- | --- | --- | --- | --- |
| Cystathionine beta-synthase | 4COO | *Sun et al. (2020)* | 0.75 | 0.68 |
| Beta-lactamase TEM | 1BTL | *Stiffler et al. (2015)* | 0.76 | 0.72 |
| Regulatory protein GAL4 | 3COQ | *Kitzman et al. (2015)* | 0.69 | 0.72 |
| Small ubiquitin-related modifier 1 | 1WYW | *Weile et al. (2017)* | 0.75 | 0.63 |
| Thiopurine S-methyltransferase | 2H11 | *Matreyek et al. (2018)* | 0.54 | 0.76 |

**Table S1.** Number of variants per class predicted by our model on the training set

| Protein | WT-like | SBI | Total loss | Low abundance, high activity |
| --- | --- | --- | --- | --- |
| NUDT15 | 1824 (68%) | 350 (13%) | 509 (19%) | 1 (<1%) |
| PTEN | 1689 (54%) | 662 (21%) | 755 (24%) | 0 (0%) |
| CYP2C9 | 2396 (57%) | 1111 (26%) | 657 (17%) | 0 (0%) |

**Table S2.** Benchmarking models generated with reduced training sets. We trained vanilla models using 1%, 10% and 50% of the set of variants in NUDT15, PTEN and CYP2C9, and evaluated them on the held-out data in these three proteins and on the independent data for the SH3 domain of GRB2 (labelled SH3). All entries are F1 scores; the first entry is relate to NUDT15, PTEN and CYP2C9 and the last two relate to the GRB2 SH3 domain.

|  | Percentage variants in training dataset |  |  |  |
| --- | --- | --- | --- | --- |
|  | 1% | 10% | 50% | 100% |
| Functional variants not used in training | 0.24 + 0.05 | 0.34 + 0.1 | 0.53+0.01 | / |
| Functional variants in SH3 | 0.15+0.1 | 0.16 +0.08 | 0.25+0.03 | 0.32 |
| All variants in SH3 | 0.40 + 0.12 | 0.61 + 0.04 | 0.62 + 0.01 | 0.63 |

**Table S3.** Benchmarking models trained on two of the three proteins. We trained vanilla models using two of the three proteins (NUDT15, PTEN and CYP2C9), and evaluated them on the held-out protein and on the independent data for the SH3 domain of GRB2 (labelled SH3). All entries are F1 scores; the first two entries relate to NUDT15, PTEN and CYP2C9 and the last four relate to the GRB2 SH3 domain.

|  | Proteins in training dataset |  |  |  |
| --- | --- | --- | --- | --- |
|  | NUDT15/CYP2C9 | NUDT15/PTEN | CYP2C9/PTEN | ALL |
| Functional variants not used in training | 0.13 | 0.33 | 0.27 | / |
| Functional residues not used in training | 0.58 | 0.51 | 0.57 | / |
| Functional variants in SH3 | 0.19 | 0.30 | 0.25 | 0.32 |
| All variants in SH3 | 0.62 | 0.62 | 0.60 | 0.63 |
| Functional sites in SH3 | 0.36 | 0.35 | 0.40 | 0.60 |
| All sites in SH3 | 0.72 | 0.71 | 0.69 | 0.75 |

**Table S4.** Enzymes analysed from the Mechanism and Catalytic Site Atlas database

| Uniprot ID | PDB | Chain | # Analysed residues | # Catalytic residues | # Functional residues |
| --- | --- | --- | --- | --- | --- |
| Q9LAK3 | 1RO7 | A | 291 | 1 | 16 |
| P82385 | 1DO6 | A | 124 | 2 | 11 |
| P09850 | 1C5H | A | 213 | 3 | 39 |
| P56868 | 1B73 | A | 254 | 6 | 17 |
| P0A7E3 | 1ORO | A | 213 | 2 | 17 |
| Q13569 | 3UFJ | A | 410 | 1 | 26 |
| P62593 | 1BTL | A | 286 | 5 | 10 |
| P00374 | 1DHF | A | 187 | 1 | 9 |
| P00469 | 1LCB | A | 316 | 3 | 25 |
| P00491 | 1RR6 | A | 289 | 3 | 16 |
| Q06241 | 1R44 | A | 202 | 5 | 11 |
| P04036 | 1ARZ | A | 273 | 2 | 25 |
| P0A6L0 | 1P1X | A | 259 | 3 | 19 |
| P19120 | 1KAZ | A | 650 | 3 | 19 |
| P77836 | 1BRW | A | 433 | 5 | 35 |
| Q16854 | 2OCP | A | 277 | 2 | 17 |
| Q53547 | 1AUO | A | 219 | 3 | 9 |
| Q55012 | 1PS1 | A | 337 | 6 | 8 |
| Q8VQN0 | 1JC5 | A | 148 | 3 | 10 |

**Table S5.** Proteins analysed from the Protein-Protein Interaction Affinity Database 2.0

| Uniprot ID | PDB | Chain | # Analysed residues | # SBI residues |
| --- | --- | --- | --- | --- |
| P0AE67 | 1FFW | A | 129 | 6 |
| P01588 | 1EER | A | 193 | 17 |
| P61972 | 1A2K | A | 127 | 19 |
| P0A0L5 | 3BZD | B | 237 | 7 |
| O77044 | 1KSD | A | 433 | 12 |

**Table S6.** Proteins used in the manuscript which are not included in the above databases

| Protein | Uniprot ID | PDB | Chain |
| --- | --- | --- | --- |
| Alkaline phosphatase PafA | Q9KJX5 | 5tj3 | A |
| Growth factor receptor-bound protein 2 (GRB2) | P62993 | AlphaFold | A |
| Disks large homolog 4 (DLH4) | P78352 | 6QJJ | A |
| Hypoxanthine-guanine phosphoribosyltransferase (HPRT1) | P00492 | 3BZD | A |
| Anti-sigma F factor | O32727 | 1L0O | A |

